# A self-limiting Sterile Insect Technique alternative for *Ceratitis capitata*

**DOI:** 10.1101/2024.12.23.629662

**Authors:** Serafima Davydova, Junru Liu, Yiran Liu, Kavya Prince, Jonathan Mann, Nikolay P. Kandul, W. Evan Braswell, Jackson Champer, Omar S. Akbari, Angela Meccariello

## Abstract

Genetic biocontrol systems have broad applications in population control of insects implicated in both disease spread and food security. In this study we establish and characterise a novel split-CRISPR/Cas9 system we term Sex Conversion Induced by CRISPR (SCIC) in *Ceratitis capitata* (the Mediterranean fruit fly), a major agricultural pest with a global distribution. Using the *white eye* gene for toolkit selection, we achieved up to 100% CRISPR/Cas9 efficiency, displaying the feasibility of *C. capitata* split-CRISPR/Cas9 systems using constitutive promoters. We then induce sex-conversion by targeting the *transformer* gene in a SCIC approach aimed for SIT-mediated releases upon radiation-based sterilisation. Knock-out of *transformer* induced partial to full female-to-male sex-conversion, with remaining individuals all being intersex and sterile. SCIC population modelling shows superior performance to traditional population control strategies, allowing for faster population elimination with fewer released sterile males. Our results build the foundation for further genetic pest control methods of *C. capitata* and related tephritid agricultural pests.

**Significance statement:** Agricultural industry faces increasing threat from a multitude of pests including the domineering tephritid fruit flies. Genetic engineering of these pests has been recently tested to develop more efficient and affordable population control strategies. Here, we develop a new approach to improve existing population control measures by testing it in one of the most famous and dangerous tephritids, the Mediterranean fruit fly. Through optimisation, we achieved desired outcomes: female fly absence achieved via semi and full female-to-male sex conversion by CRISPR-mediated genome editing through gene mutations. For the first time in this insect, we used a split, and thus inducible, approach for such genome editing. Our work holds the potential to significantly improve tephritid population control strategies.

## Introduction

Insect population management stands to greatly benefit from the application of endogenous CRISPR/Cas9 tools (1–2). CRISPR/Cas9-mediated gene editing in several disease vector and agricultural pest species has now been implemented via both ribonucleoprotein delivery and endogenous expression of its components (3–6). Single cassette-based expression has been used for the establishment of genetic control systems such as homing gene drive (7–9) and X-chromosome shredding or poisoning (10–11). More safeguarded split gene drives also exist, whereby gRNA and Cas9 are expressed from separate loci (9, 12–14). These technologies aim to replace current population control strategies including the sterile male production-dependent Sterile Insect Technique (SIT) (15).

To test CRISPR/Cas9 activity *in vivo* in both model and non-model insects, essential efforts have also been made through establishment of split CRISPR/Cas9-expressing lines (4, 16). Due to their ‘inducible’ nature, binary CRISPR/Cas9 systems have broader implications in research of insects with immense economic significance, including the ability to accessibly uncover gene functions *in vivo*. Such work will accelerate scientific progress, meeting the demand for increasingly looming global spread of insect disease vectors and agricultural pests. Another application of split-CRISPR/Cas9 is precision-guided SIT (pgSIT) developed in *Drosophila melanogaster*, and thereupon tested in *D. suzukii* and mosquito disease vectors (17–21). Dissimilarly to other existing single and split-cassette strategies, pgSIT simultaneously addresses two major limitations of SIT: manual sex-separation requirement and reduced male fitness associated with irradiation-based sterilisation (22). In line with the primary principle of SIT, and in contrast with homing gene drives, pgSIT is self-limiting in nature as its final product consists of entirely sterile males. This is achieved via crosses of Cas9-expressing and gRNA-expressing lines together, which in turn induces simultaneous knock-outs of a male fertility and a female development-implicated genes (18).

For the past half century, SIT has been increasingly implemented to prevent population growth of the abundant and invasive tephritid agricultural pests (23). It has been notably efficient for the management of the Mediterranean fruit fly (medfly), *Ceratitis capitata,* a vastly polyphagous tephritid pest with an existing worldwide prevalence (24–25). Tephritid SIT efficiency can be boosted using traditional genetic sex-sorting systems (26–29), although they suffer from associated recombination-dependent genetic instability and fertility problems (30). While transgenic sexing alternatives have been tested (31–35), none have been deployed on a frequent basis in SIT programmes. More stringent and efficient measures of population control are therefore still necessary to oversee the numerous tephritid migrations, which are currently benefiting from the rising temperatures and globalisation (36–38). Using CRISPR/Cas9-mediated approaches may consequently offer a novel solution to tephritid population management.

Here, we build the basis for a future establishment of self-limiting pgSIT in *C. capitata* and related tephritids by proxy. Thus far, we endogenously co-expressed gRNA and Cas9 in the medfly in single cassette systems whereby Cas9 expression was purposefully limited to the germline (11, 39). Since no split CRISPR/Cas9 system exists in the species to date, we sought to initially select an appropriate CRISPR/Cas9 toolkit by evaluating germline and constituent promoters for Cas9 expression, in addition to testing a new U6-modified promoter for gRNA expression. For this, we implemented a mixture of characterised and uncharacterised regulatory elements and targeted the *white eye* gene with 2 gRNAs at once. We also pursued the knock-out the *transformer* (*tra*) gene alongside a *ß-tubulin* (*ß-tub*) homologue, hoping to interfere with female development and male fertility accordingly. As *tra* knock-out induces partial and complete female-to-male sex-conversion in the medfly (39–40), we anticipated that females would be transformed into intersexes and males, thus doubling the output of our system. This resulted in the generation of a novel self-limiting and genetic biocontrol system we term Sex Conversion Induced by CRISPR (SCIC) which acts by dominantly converting females into males which can be used for population control. To understand the dynamics of SCIC, we used simulations to model the release of hereby generated flies and hence make a comparison with traditional SIT.

## Results

### CRISPR/Cas9 toolkit selection using the *white eye* gene

To establish a binary CRISPR/Cas9 system, two transgenic *C. capitata* strains are required, separately expressing Cas9 and at least two different, double gRNA(dgRNA). Prior to tackling genes involved in female development and male fertility, we aimed to optimise the *C. capitata* CRISPR/Cas9 toolkit via targeting the *white eye* gene (GeneID_101458180) (41–42), the subject of extensive CRISPR/Cas9 testing previously (6, 11, 39, 43). For this purpose, two types of *piggyBac* constructs were generated: one containing Cas9 and the other containing double-guide RNA (dgRNA) targeting *white eye* using two newly selected gRNAs under an endogenous U6 promoter (GeneID_LOC111591841). This U6 promoter has an additional 71 upstream base pairs and a 159 base pair deletion in its sequence compared to its longer version that has been tested for gRNA expression previously (11) Regulatory elements from three genes were assessed for Cas9 expression: namely *D. melanogaster polyubiquitin* (GeneID_38456) (44), endogenous *polyubiquitin* (GeneID_101461787) and endogenous *nanos* (GeneID_101451248) (45). The dgRNA construct included a *Hr5-IE1-eGFP* marker, whilst Cas9 constructs were uniformly marked with *Hr5-IE1-DsRed.* To visually verify Cas9 expression, a T2A peptide was additionally used to link *Cas9* and the downstream *GFP*. All constructs were delivered into wild-type Benakeion *C. capitata* embryos for germline transformation. Inverse PCR was used to identify independent *piggyBac* integrations in fluorescent marker-positive G1 flies (Supplementary Table 1; Supplementary Figure 1). Altogether, four unique strains were characterised for the *nanos*-Cas9 (CcNos.1-4) and *D. melanogaster polyubiquitin*-Cas9 constructs (DmPub.1-4); and one strain each was established for the dgRNA (We.1) and the *endogenous polyubiquitin*-Cas9 (CcPub.1) constructs. All but one generated stain achieved homozygosity. We concluded that the DmPub.4 strain had lethality among individuals with two copies of the cassette as crosses between heterozygous individuals never resulted in homozygote occurrence.

We proceeded to assess the suitability of the CcNos, DmPub and CcPub strains for the designed split CRISPR/Cas9 system. Cas9-harboring females from each strain were crossed with dgRNA-harbouring (We.1) males and their F1 progeny was analysed for eye colour. In the crosses with CcNos.1-4, all observed F1 flies were universally red-eyed. Trans-heterozygous DsRed+/GFP+ females from each of the four crosses were mated with males form a homozygous recessive *white eye* mutant strain, generated in previous work (11). The resulting F2 white eye phenotype frequency ranged from 0.0 to 12.5% across the four strains (CcNos.1-4) with no mosaism observed (Supplementary Figure 2), altogether indicating insufficient Cas9 expression with the *nanos* promoter. Chi-squared tests identified no statistically significant differences from the expected 100% red eye phenotype for any of the CcNos.1-4 strains. To validate that Cas9 was expressed in the ovaries of Cas9-harboring females, mature females were dissected and checked for GFP fluorescence. Strong GFP signal was detected in both tested CcNos strains (CcNos.3-4) (Supplementary Figure 3).

In the initial crosses with *D. melanogaster polyubiquitin*-Cas9 females, the F1 progeny largely consisted of non-red eyed individuals, indicative of the presence of biallelic knock-outs. We observed red, mosaic, orange, and white eye phenotypes (Supplementary Figure 2). Non-red eyed fly frequency ranged from 63.1 to 93.5% across the DmPub.1-4 crosses with We.1. Chi-squared tests indicated statistical significance (p < 0.0001) for each of the four strains. Briefly, we performed molecular analysis of *white eye* target sites of F1 DsRed+/GFP+ flies with different eye colours, selected at random across the four crosses. We detected a fragment with a complete deletion of ±252 bp between the two gRNA target sites in all tested individuals, confirming high Cas9 activity. We set up crosses for DsRed+/GFP+ red-eyed F1 females for DmPub.1 and DmPub.3 strains with *white eye* mutant males. The resulting F2 progeny had 57.0% and 42.9% non-red eye phenotypes for DmPub.1 and DmPub.3 respectively, elusive of monoallelic pUb-Cas9 activity at F1 (Supplementary Figure 2). Simultaneously, no DsRed+/GFP+ red-eyed flies were observed among the DmPub.2 and DmPub.4 F1 individuals. We therefore deduced that for these strains all flies with both Cas9 and dgRNA components were non-red eyed indicative of complete CRISPR/Cas9 efficiency. Repeat crosses were set up between DmPub.2 and DmPub.4 females, with homozygous males from the We.1 dgRNA strain, and F1 flies were additionally screened for DsRed and GFP fluorescent markers. The DsRed+/GFP+ F1 medflies had 100 % non-red eye phenotypes in both crosses (Figure 1). Importantly, all DsRed-/GFP+ DmPub.4 F1 offspring still exclusively had only white and mosaic phenotypes, suggestive of strong maternal Cas9 carryover (Supplementary Table 2).

**Figure 1.**
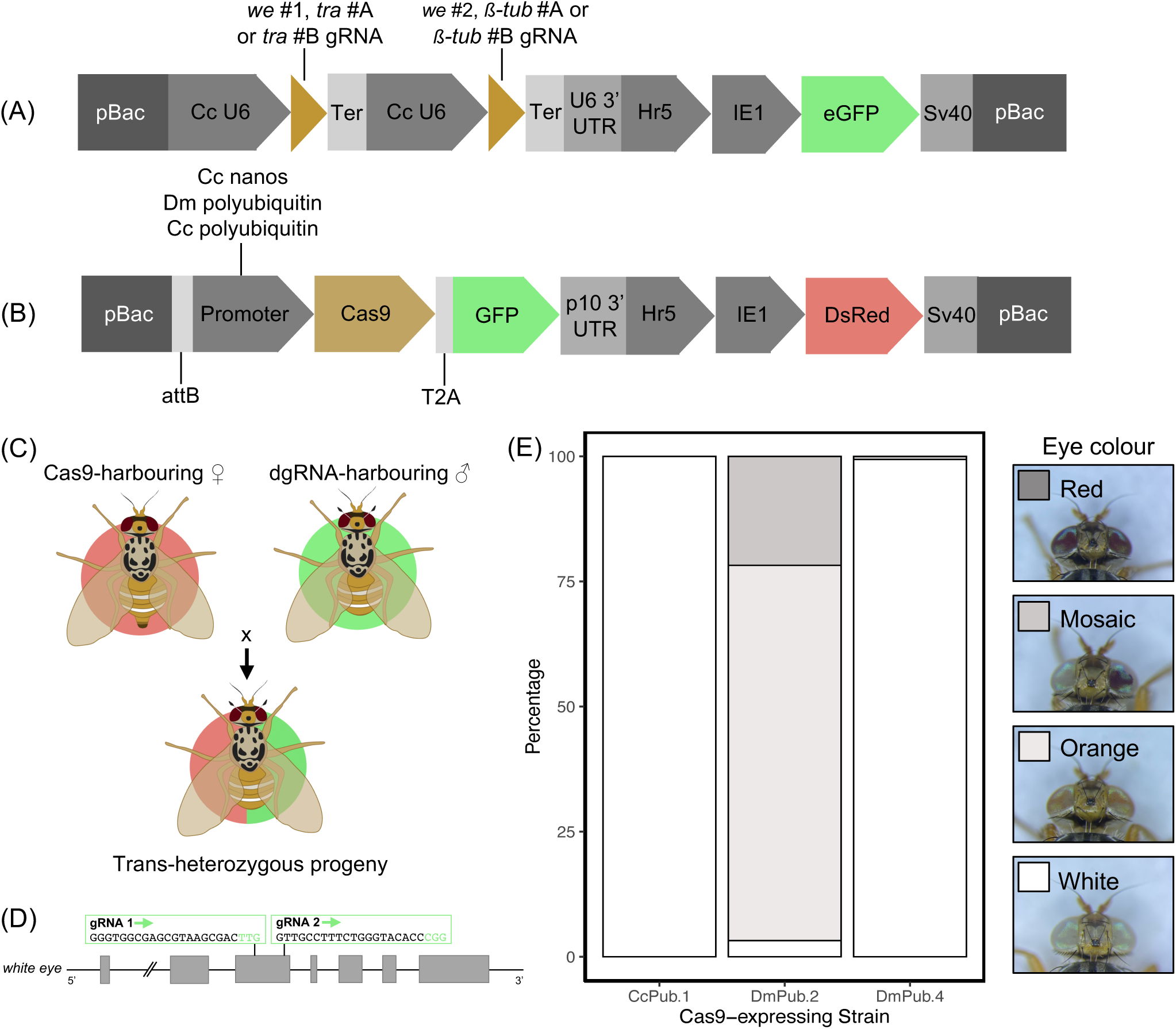
Split CRISPR/Cas9 system design and optimisation against the *white eye* gene. Schematic representations of (A) double-guide RNA (dgRNA)-harbouring and (B) Cas9-harbouring constructs used for generation of *Ceratitis capitata* transgenic lines. (A) The dgRNA, all under the control of an endogenous U6 promoter, were either targeting the *white eye* gene, or simultaneously targeting *transformer* and *ß-tubulin* genes. (B) The Cas9 constructs varied by promoter sequences, whereby endogenous *polyubiquitin* and *nanos*, and *Drosophila melanogaster polyubiquitin* were used. Cas9 was uniformly linked to GFP via a T2A peptide. (C) A schematic representation of the system design, whereby Cas9 and dgRNA expressing lines were crossed together (F0), and their F1 progeny was assessed for phenotypic outcomes. (D) A diagram indicating the positions of gRNAs within the *white eye* gene with both targets located on exon 3. (E) A stack graph showing percentage phenotypes of the DsRed+/GFP+ F1 progeny of crosses between Cas9-harbouring females with dgRNA-harbouring males from the We.1 strain. No red-eyed F1 flies were observed among the progeny of the three Cas9-expressing strains shown. (E) was constructed in RStudio (*ß-tub*, *beta-tubulin*; Cc, *Ceratitis capitata*; Dm, *Drosophila melanogaster*; Ter, terminator; *tra*, *transformer*; *we*, *white eye*).

Endogenous *polyubiquitin*-Cas9 (CcPub.1)-harbouring females were also crossed with dgRNA (We.1)-harbouring males. DsRed+/GFP+ and DsRed-/GFP+ flies universally exhibited a white eye phenotype (Figure 1; Supplementary Table 2). As with its exogenous counterpart, the molecular analysis highlighted the presence of a complete deletion between the two gRNA target sites in the *white eye* gene. Additionally, ovaries of females of both DmPub.4 and CcPub.2 strains exhibited GFP fluorescence (Supplementary Figure 3). Altogether, these experiments unveiled the suitability of three Cas9-harboring strains (DmPub.2; DmPub.4; CcPub.1) for further testing.

### Achieving sex-conversion by targeting *transformer* in a *C. capitata* split CRISPR/Cas9 system

To test the fully fledged SCIC system in *C. capitata*, we used two new *Hr5-IE1-eGFP*-marked dgRNA constructs. They contained gRNAs now targeting *tra* (GeneID_101456163) and a previously untested homologue of the *D. melanogaster ß-tub* gene (GeneID_101453087), which aimed to induce female-to-male sex-conversion and male sterility, respectively. The constructs, entitled GuideA and GuideB, differed amongst each other by the gRNA sequences only, with GuideB targets located further downstream within the coding sequences of both genes (Figure 2). In this way, we sought to test the optimal target sites for Cas9-mediated knock-out. We thereupon established and characterised two independent strains for each construct via inverse PCR (GuideA.1-2; GuideB.1-2) (Supplementary Table 1; Supplementary Figure 1).

**Figure 2.**
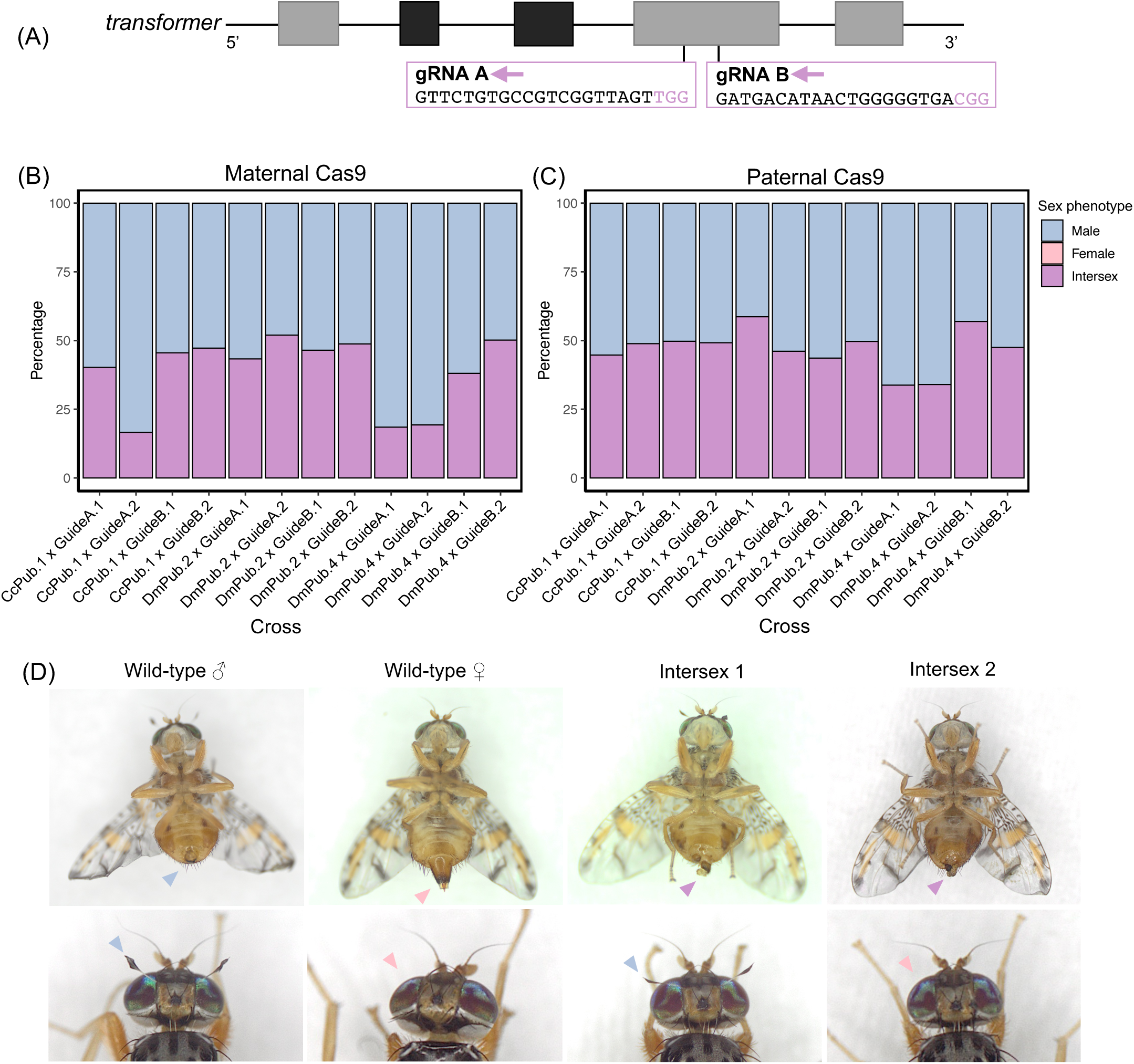
Split CRISPR/Cas9 system activity. (A) A simplified diagram showing the positions of gRNA targets within the *transformer* gene. The universal exons are shown in light grey, whilst the male-specific exons containing premature stop codons are shown in dark grey. (B, C) Stack graphs showing sex phenotype distributions of trans-heterozygous F1s resulting from crosses between Cas9-expressing strains (DmPub.2; DmPub.4; CcPub.1) and dgRNA-expressing strains (GuideA.1-2; GuideB.1-2) whereby the Cas9 was supplied maternally (B) (n = 2,581) or paternally (C) (n = 1,966). No females were found among the 4,577 DsRed+/GFP+ flies screened across the 24 cross types. (D) Representative images of newly-eclosed F1 XX intersexes alongside wild type male and female controls. Male features (genitalia and orbital bristles) are indicated with blue arrow heads; female features (ovipositor and absence of orbital bristles) are shown with pink arrow heads; and intermediate intersex-unique genitalia are highlighted with purple arrow heads.

Next, males from the GuideA.1-2 and GuideB.1-2 strains were crossed with females from pre-selected Cas9-expressing strains (DmPub.2; DmPub.4; CcPub.1). The resulting F1 DsRed+/GFP+ populations lacked females, consisting entirely of males and intersexes (Figure 2B-C). In the crosses with DmPub.4 and CcPub.1 Cas9-expressing strains, all DsRed-/GFP+ F1 flies were also exclusively male or intersex, indicative of maternal Cas9 carryover (Supplementary Table 3). However, when reciprocal parental crosses were carried out with Cas9-expressing males, DsRed-/GFP+ populations were comprised of males and females instead (Supplementary Table 3). Additionally, in the crosses with male Cas9-expressing parents, male sex bias only reached a maximum of 66.2% compared to an averaged 83.5% maximum observed with Cas9-expressing mothers (Figure 2B).

### Characterisation of *transformer* knock-out in F1 flies

Amongst the 12 crosses between Cas9 strains (DmPub.2; DmPub.4; CcPub.1) and dgRNA strains (GuideA.1-2; GuideB.1-2), the observed intersexes exhibited male-like secondary sex characteristics suggestive of elevated levels of sex-conversion and thus elevated Cas9 activity (Figure 2D). To investigate the morphology of their internal genitalia, sexually mature intersexes were dissected, pooled together by crosses of the three Cas9 strains (DmPub.2; DmPub.4; CcPub.1) with either GuideA or GuideB-harbouring dgRNA strains. The phenotypes ranged from two fully formed ovaries to two fully formed testes, with 12 individuals having no internal genitalia at all (n = 30) (Supplementary Table 4). Additionally, throughout the experiment intersexes laid no eggs, and upon dissection their ovaries were visibly larger than those from wild-type females, further eluding to their inability to oviposit eggs.

To explore the potential differences in gRNA cleavage among the two gRNAs, *tra* target PCR was completed and sequenced on both male and intersex DsRed+/GFP+ F1s, with a karyotyping PCR performed to verify their corresponding sex-chromosome profiles. Across 179 successfully sequenced flies, sampled from all 12 crosses (DmPub.2; DmPub.4; CcPub.1 x GuideA.1-2; GuideB.1-2) 178 possessed mosaic residues around the gRNA sequences in the genome. Upon sequence deconvolution, cleavage rates were estimated to be as high as 100% with indel mutations largely consisting of short deletions (Supplementary Figure 4). Collectively across the 12 crosses (DmPub.2; DmPub.4; CcPub.1 x GuideA.1-2; GuideB.1-2), XX males were uncovered amongst phenotypic males (9.2%, n = 119). To further understand the ratio of sex-converted XX males in the populations, larger-scale karyotyping PCRs were conducted on F1 DsRed+/GFP+ males from two selected crosses. Specifically, DmPub.2 x GuideA.2 was chosen as the cross that adhered to 1:1 intersex: male ratios, whilst CcPub.1 x GuideA.2 cross was investigated due to its strong male bias (83.5%) (Figure 2). Overall, 1.1% (n = 92) and 13.0% (n = 100) of males had XX karyotypes in F1 progeny of DmPub.2 x GuideA.2 and CcPub.1 x GuideA.2 crosses accordingly, indicating the presence of fully female-to-male converted XX individuals.

To evaluate the potential fitness costs of the F1 trans-heterozygous progeny, the F1 egg-adult survival was set-up whereby five parameters were assessed: laid egg count, egg hatching rate, hatched larval-pupal recovery, pupal-adult recovery, and total egg-adult recovery (Supplementary Figure 5). Hereby, Cas9-expressing (DmPub.2; DmPub.4; CcPub.1) females were crossed with gRNA-expressing (GuideA.1-2; GuideB.1-2) males, alongside wild-type controls. Kruskal-Wallis and Dunn multiple comparisons tests were used for statistical analyses. Seven out of 12 experimental crosses had a statistically significant decrease in egg-adult survival from wild-type. This was particularly notable for all crosses with CcPub.1 strain (Supplementary Figure 5). Two crosses (DmPub.2 x GuideA.2; CcPub.1 x GuideA.2) were selected based on their opposing sex-conversion rates recorded previously (Figure 2), to verify whether XX embryos persisted until adulthood. This was achieved through the repetition of F1 egg-adult assays and consequent adult phenotype scoring (Supplementary Figure 6). In spite of lower sex-conversion rates amongst the F1 progeny in the CcPub.1 crosses in this experiment, the egg-adult survival did not exceed initial experiments that lacked adult phenotype scoring, which may be due to variable Cas9 expression or female lethality at embryonic stages of development. To further examine these hypotheses, we also established line egg-adult assays. Both sibling crosses and crosses with wild type flies which aimed to mimic two copy and one copy transgene inheritance respectively (Supplementary Table 5). There were notable reductions in survivorship across the Cas9-expressing strains, and the largest egg-adult decrease was observed in the CcPub.1 strain (Supplementary Figure 7). These results connote that the observed lower F1 egg-adult survival in select crosses could be attributed to the fitness costs associated with harbouring a Cas9-expressing cassette, rather than F1 female lethality.

### Assessing the *ß-tub* homologue function in *C. capitata* fertility

To establish a complete pgSIT system, targeting a gene implicated in male fertility was required, alongside targeting *tra*. *ß-tub* was therefore selected as a target due to its well-characterised involvement in spermatogenesis in *D. melanogaster* (46). We verified CDS sequence of the *D. melanogaster* gene (GeneID_CG9359) with the *C. capitata* genome and uncovered a predicted gene with 73.9% similarity (GeneID_101453087) (Supplementary Figure 8). Due to inclusion of its gRNAs in GuideA and GuideB constructs, we expected that its knock-out would result in fertility decline in DsRed+/GFP+ F1 males. Trans-heterozygous males were thus crossed with wild-type females and egg laying and egg hatching rates were measured as measures of fecundity and fertility, respectively. Statistically significant reductions compared to wild type were only observed in egg hatching rates of flies from three out of 12 grand-parental crosses (Supplementary Figure 9). The trends of egg hatching rates in flies with CcPub.1 grandparents resembled those observed during earlier line and F1 egg-adult assays. Overall, the lack of consistent and complete reductions in egg hatching rates implied that knocking-out the selected *ß-tub* target gene was not sufficient to induce male sterility. To eliminate the possibility that neither gRNA was efficient, a target PCR was performed on the same males and intersexes as utilized for *tra* cleavage verification. PCR products sequenced whereby highly mosaic residues were observed around the gRNA sequences. Similarly to *tra*, both gRNAs were successfully cleaved up to a predicted 100% in males and intersexes alike (Supplementary Figure 4). This data suggests that despite successful disruption of *ß-tub*, it cannot be used as a singular target to tackle male fertility, which may be attributed to redundancy in its function or inappropriate gene selection.

### Models show suppression by SCIC and pgSIT releases in *Ceratitis capitata*

To show the possible performance and time scale for SIT methods in *Ceratitis capitata*, we constructed a model with weekly time steps and examined the effects of releasing SIT males into a panmictic *C. capitata* population. We included standard radiation-based SIT, in which the release ratio represented the ratio of the number of sterile males released per week to the number of wild-type males when the population is at equilibrium. We also included variants based on GuideA.2 performance, either with DmPub2 or CcPub1 from the SCIC system. These could be sterilised by radiation with attendant male mating success fitness costs, or alternatively, sterilised by hypothetical use of additional sgRNAs targeting a male fertility gene (pgSIT), which we assume has no fitness cost. Based on our experimental data for an equivalent amount of food in a rearing facility, the DmPub2 line is expected to have the same survival as wild-type, but benefits from a small fraction of XX males, making the effective release size 1.1% higher. The CcPub1 line has only 85% survival, but benefits from 35% of XX individuals developing as males, giving this system as effective release size 15% higher than standard SIT. We pessimistically assumed based on first-male mating preference that a subsequent mating with wild-type would produce 50% fertility, and that fertility would remain at 100% if the first male was wild-type.

With a modest weekly release ratio of 0.5, no system could induce population suppression, and with a ratio of 2, all five systems functioned well (Figure 3A). However, when the release ratio was 1, only the CcPub1-based pgSIT system was able to eliminate the population within 100 weeks (though the radiation-based SCIC system with CcPub1 came close), reducing it to below 90% after just 75 weeks. The DmPub system enabled substantial population reduction in pgSIT form, but the standard SIT release program and the DmPub system combined with radiation-sterilisation gave similar performance, only moderately reducing the population. Systematically evaluating the effect of varying the release ratio (Figure 3B), it is clear that substantial benefits in improving the time to population elimination are accrued until the release ratio reaches 3. Afterward, higher efforts are needed to reduce the time for population elimination. When release sizes are insufficient to eliminate the population, the number of adult females can still be substantially reduced after 100 weeks of releases (Figure 3C).

**Figure 3.**
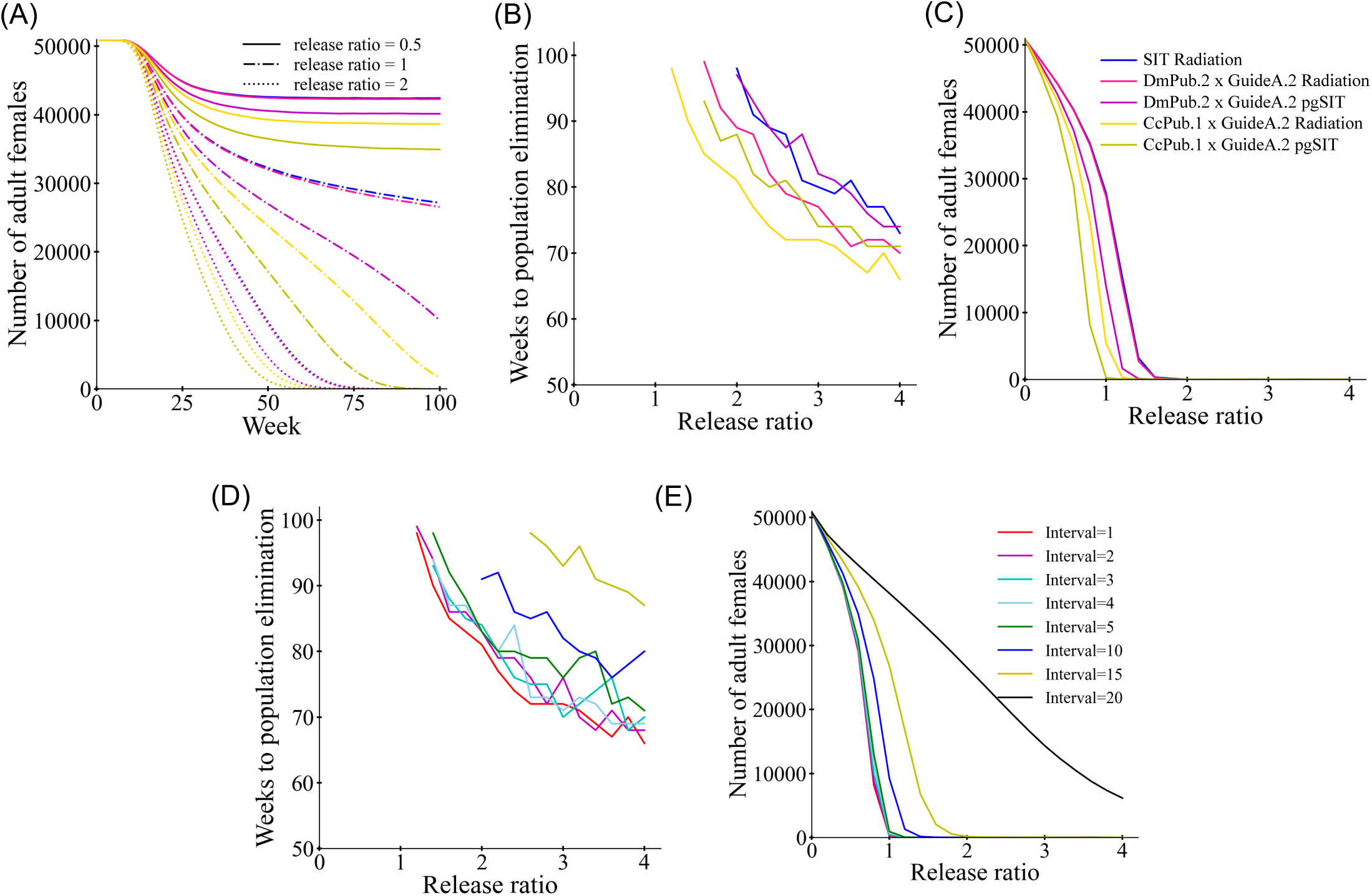
**Modeling SIT in *Ceratitis capitata*** SIT males were released into a population of medflies (with 50,000 adult females) with a variable effort represented by the release ratio. This represents the number of standard SIT males released per week as a fraction of the normal adult male population, and an equivalent effort for SCIC-radiation and pgSIT-based methods on the same amount of food in the rearing area. (A) The female population through time for each of three SIT techniques with three release ratios. The (B) time to population elimination or (C) average number of females during the last 20 weeks with varying release ratio for each system with weekly releases. Additional simulations with varying release interval for CcPub.1 x GuideA.2 showing (D) time to population elimination or (E) average number of females during the last 20 weeks. The release interval represents the weeks between releases (the total release number for the whole time period remains the same, so large numbers mean fewer but larger releases.

Because medflies can be relatively long-lived, we examined the possibility of extending the interval between releases. This can make rearing them in batches simpler if there are fewer overlapping batches being reared at any one time point. We found that if there are 10 or more weeks between releases, suppression efficiency substantially drops (Figure 3D-E). However, with an interval of 4 or less, suppression efficiency appeared nearly the same as with weekly release intervals.

## Discussion

We report on the establishment and characterisation of a highly efficient split CRISPR/Cas9 system in *C. capitata* as a precursor for further application in technologies such as pgSIT. First, the limited single cassette CRISPR/Cas9 toolkit in the medfly (11, 39) was expanded when three Cas9 (endogenous *nanos*, *D. melanogaster polyubiquitin*, endogenous *polyubiquitin*) and one endogenous U6 gRNA regulatory regions were assessed by targeting the *white eye* gene. By novelly using constituent endogenous and exogenous *polyubiquitin* promoters for Cas9 expression, we were able to achieve desired phenotypes in up to 100% trans-heterozygous progeny when gRNA and Cas9-expressing lines were crossed together. Due to observed optimal efficiency, selected strains, characterised in this work, can be used to accelerate downstream medfly CRISPR/Cas9 experiments such gRNA and Cas9 promoter selection. In the F1 trans-heterozygotes, the notable differences in offspring phenotypes can be attributed to integration-dependent variation in Cas9 expression, exemplified by the orange, mosaic and white eye phenotypes in *D. melanogaster polyubiquitin*-Cas9 crosses (Figure 1). Unexpectedly for both *polyubiquitin* promoters, abundant maternal Cas9 carryover was detected, which was similarly dependent on the genomic context of the Cas9 cassette integration sites (Supplementary Table 2).

To generate a sex-conversion-based system we call SCIC, we also constructed and tested gRNA-expressing lines targeting the *C. capitata tra* gene. A plethora of studies have shown that knock-out or knock-down of *Tephritidae tra* or its accompanying gene *transformer-2* results in XX embryo sex-conversion into intersexes or males (39–40, 47–51). Expectedly, we obtained mixed populations of males and intersexes when Cas9 and gRNA-expressing lines were mated together. Across all performed crosses, for the first time no Cas9+/gRNA+ females were observed, supportive of elevated Cas9 activity seen in preliminary *white eye*- targeting crosses. With molecular verification we concluded that XX embryos were undergoing partial and full sex conversion into phenotypic intersexes and males respectively. Even though a large proportion of XX flies is not fully converted into males, the resulting sterile intersexes cannot damage hosts directly due to ovipositor absence in 100% of individuals. All F1 flies can thus be sterilised and distributed into the wild without a need to sex-separation. This would allow for more cost-effective SIT programmes by doubling released populations and preventing preferential mating of co-released males and females.

The witnessed high Cas9 expression, however, had a notable effect on fly fitness, particularly in the endogenous *polyubiquitin*-Cas9-harboring strain (Supplementary Figure 7). Moving forward, site-specific integrations may benefit with selection of Cas9-expressing lines which balance high fitness with sufficient endonuclease activity, similarly to the DmPub.2 strain generated herein. This can be achieved using established homology-directed-repair-dependent methodologies (39, 47, 52) or non-homologous end-joining alternatives (53), not yet tested in tephritids.

Alongside the well-performing constituent promoters, we sought to investigate the capability of germline candidates to produce desired outcomes in a split CRISPR/Cas9 system in *C. capitata.* We anticipated that the truncated version of the characterised *C. capitata nanos* promoter (39) would provide adequate Cas9 expression necessary for *white eye* mutagenesis. However, *white eye* was cleaved insufficiently despite the high GFP signal observed in ovaries of *nanos*-Cas9-expressing females. Spaciotemporal differences of expression patterns of CRIPSR/Cas9 elements in the gRNA and four Cas9 strains may therefore be underpinning low target cleavage rates. To assess whether germline promoters can be employed in our split CRISPR/Cas9 *C. capitata* system, the widely used *vasa* and *zpg* could be used next (18, 20, 39, 54).

Targeting *ß-tub* alongside *tra* proved ineffective in inducing male sterility. We detected fertility reductions which were later attributed to parental line fitness costs resulting from transgene cassette expression. As both tested gRNAs targets were cleaved with extreme efficiency, the selected *ß-tub* gene needs to be explored further to determine whether it was appropriately chosen or whether its function can be compensated by other genes. To establish a functional pgSIT system, other spermatogenesis-implicated genes need to be selected in its stead, such as the characterised testis-restricted *ß2-tubulin* (55–56). Genes expressed in both male and female germlines may offer an additional alternative which includes *innexin-5*, the knockdown of which produced sperm-less males and egg-less females in *C. capitata* (57). This approach is possible for the use in our system as internal genitalia formation in intersexes is already affected by *tra* knock-out (Supplementary Table 4) and is altogether irrelevant due to their infertility.

Our model showed the increased efficiency of the SIT releases when targeting *tra* due to formation of fertile XX males, even at the cost of lower viability. Timescales for suppression with moderate release ratios were under two years, though the population was substantially reduced after just a year of releases. In more complicated ecological environments with seasonality and competing species, it may be possible to reduce the population even more quickly with appropriate release timing. It should be noted that our model has several simplifications that may affect considerations for SIT programs. First, we did not model any fitness cost of released males from being reared in a facility compared to males that developed in the wild. This is likely to affect all systems equally, though it can potentially be mitigated by pre-release aromatic treatment (58). Second, it remains unclear if XX *tra* knockout males retain the same mating success as wild-type males, which could reduce the efficiency of the CcPub1 system. Next, the timescale of suppression is highly dependent on the generation time. Our age-based mortality can vary between strains, but mortality from other sources is likely to also be very important. Our 10% weekly mortality from other sources was an estimate, and higher values could speed population elimination and reduce weekly release ratios, though closer release intervals would also be required in this case. Finally, the exact nature of density-dependence can increase or decrease suppression difficulty. Nevertheless, with the success of standard SIT programs in the medfly, the prospects for successful genetic SIT deployment are good if the costs for maintaining and crossing the Cas9 and gRNA strains is not high.

For cost reductions in SIT, a plethora of sex-sorting strategies have been developed in tephritid species via both traditional (29–30), and more recently, genetic engineering-based approaches (31–32, 34–35). Although sex-sorting is not a requirement for the F1 trans-heterozygotes of our system, their parents need to be selected by sex. As females need to be preserved until mating, early female lethality technologies, such as those relying on tetracycline are unapplicable (32; 34). Thus, to alleviate the labour-intensive task of sex separation, our system can be further coupled with a cisgenic or transgenic fluorescent marker-reliant sex-sorting strategy aimed to rely on automated separation exclusively (31; 59).

Whilst SIT-based population control of *C. capitata* continues to be improved, its efficiency in related tephritid pest species has not resulted in the same level of success. Among novel genetic approaches, sex-distortion systems have been reported in the medfly (11, 39), however neither combine female-to-male conversion with self-limiting characteristics. Our work fills this gap using *C. capitata* as a model for other tephritid species, which lack extensive CRISPR/Cas9 and sex-ratio manipulation tools. This study lays the groundwork for accessible construction of tephritid pgSIT systems with the goal of self-limiting, sustainable and cost-effective population control.

## Methods

### Promoter and target selection and molecular cloning

All plasmids were constructed using a preexisting *piggyBac* plasmid. Specifically, a construct containing two U6 promoters with terminators was created using GeneBlock 1141A-1 and 1141A-2. Two gRNAs per construct were amplified via PCR using the primers listed in Supplementary Table 6 and inserted into the U6-containing plasmid after linearising it with ApaI. Two gRNAs were selected to target each gene using the CHOPCHOP tool (60). For the *white-eye* gene (GeneID_101458180), both gRNAs target the third exon, following the rationale previously described (43). Notably, gRNA #WE3 specifically targets the ABC transporter signature sequence. For the *tra* gene (GeneID_101456163), two gRNAs targeting the female-specific *Cctra* exon were selected; and for *ß-tub* (GeneID_101453087), two gRNAs targeting exon 2 were chosen.

For all Cas9 constructs, a preexisting plasmid containing Cas9-T2A-GFP was linearised using PmeI and XhoI. The putative nanos promoter (GeneID_101451248) includes a 611 bp sequence upstream of the 5’ UTR, followed by a 163 bp 5’ UTR region. The putative polyubiquitin-C promoter (GeneID_101461787) consists of an 1849 bp sequence upstream of the polyubiquitin-C coding region. The promoters were first amplified using the primers listed in Supplementary Table 6 and inserted into the plasmid via Gibson assembly.

### C. capitata husbandry

The *C. capitata* WT Benakeion and transgenic strains were reared as previously described (11). The homozygous recessive *we-/-* strain with a mutation in exon 3 of the *white eye* gene, used in this study, was generated, and characterized in the lab previously (11). Two recipes of fly food were used to sustain the larvae (43; 61), whilst adults were consistently fed a mixture of yeast and sugar in equal proportions. All fluorescence screening and image enquiry was completed using the MVX10 Macro Zoom Fluorescence Microscope System (Olympus).

### Microinjections and transgenic line establishment

To induce germline transformation, microinjections of all abovementioned constructs were performed into WT Benakeion embryos. The *piggyBac* plasmids were delivered alongside a transposase-containing helper (62) in accordance with conditions described previously (11). G0 adults were reciprocally crossed to WT Benakeion flies. G1 progeny was screened for fluorescence and up to 10 marker-positive flies were individually crossed to 10 WT Benakeion flies of the opposite sex. Upon establishment, all transgenic lines were maintained via sibling crosses of 10-15 males with 20-25 females.

### Inverse PCR

The *piggyBac* construct integrations were analysed in positive G1 flies post-mating. Genomic DNA (gDNA) from whole flies was extracted using an adapted phenol-based methodology characterized previously (63). To identify *piggyBac*-neighbouring genomic sequences in the extracted gDNA, inverse PCR was performed as summarised before (11). The final PCR products were run on 1% agarose gel and Sanger sequenced (Genewiz Inc.). The latest *C. capitata* genome re-assembly (GenBank GCA_905071925.1) was used for sequence analysis.

### Test crosses between Cas9 and gRNA-harbouring strains

For the *white eye* gRNA tests, homozygous gRNA (We.1)-expressing males and Cas9 (CcNos.1-4; DmPub.1-4; CcPub.1)-expressing females were crossed together to obtain F1 gRNA+/Cas9+ trans-heterozygotes which were screened for eye colour and classified as ‘white’, ‘orange’, ‘mosaic’ or ‘red’. After the initial *white eye* crosses of all Cas9-harbouring strains (CcNos.1-4; DmPub.1-4), additional CcPub.1, DmPub.2 and DmPub.4 crosses were carried out, and the resulting F1 flies were additionally screened for DsRed and GFP fluorescence markers. For the DmPub.4 strain, the heterozygous individuals were crossed instead as no homozygous flies survived to adulthood. If present, the red-eyed DsRed+/GFP+ F1 flies were crossed with the *white eye* mutant *(we* -/-) Benakeion strain to generate F2 progeny from which the frequency of ‘CRISPANT’s was derived. For the *tra/b2-tub* gRNA tests, gRNA (GuideA.1-2; GuideB.1-2) and Cas9 (DmPub.2,4; CcPub.1) stains were crossed together in a reciprocal fashion. The resulting F1 progeny was screened for DsRed, GFP and sex phenotypes, which were categorized as ‘male’, ‘intersex’ or ‘female.’

### Molecular characterization of the *we* target

Up to 10 F1 trans-heterozygotes of each observed eye colour were randomly collected for molecular analysis. Their gDNA was used as template for PCR amplification of the *white eye* exon 3 fragment containing both gRNA target sites with primers designed in Geneious Prime 2023.1.2 (Supplementary Table 6). PCR products of interest were purified for Sanger sequencing (Genewiz Inc.) either via gel extraction with Monarch® DNA Gel Extraction Kit (New England Biolabs® Inc.) or directly after PCR using the Monarch® PCR & DNA Cleanup Kit (New England Biolabs® Inc.).

### Fitness assays

To assess the fitness of transgenic Cas9-expressing and gRNA-expressing strains egg-adult assays for selected strains were carried out in technical triplicates using 10 male and 20 female flies. All performed crosses are summarised in Supplementary Table 5. 5-hour eggs collections and hatching rate determinations were performed as described previously (31) using Fiji (64) for manual egg counting. Pupal and adult recovery rates were recorded thereafter. To determine F1 trans-heterozygote survival, identical experiments were also established for gRNA x Cas9 crosses in six biological replicates using 5 gRNA-harbouring males and 15 Cas9-harbouring females. Later, DmPub.2 x GuideA.2 and CcPub.1 x GuideA.2 crosses were repeated in biological triplicates using 10 males and 20 females. The resulting adults were also screened for GFP, DsRed and sex phenotypes.

### Abdominal dissections

On the day of eclosion, DsRed+/GFP+ intersexes were transferred to cages and reared under standard conditions for five days to achieve full sexual maturity. The abdominal dissections were conducted using a surgical needle on sterile glass slides in phosphate buffered saline (PBS). Similar dissections were performed for Cas9-expressing females, the ovaries of which were additionally imaged using RFP and GFP filters.

### F1 fertility assays

F1 males were subjected to further fertility analysis via crosses of 10 DsRed+/GFP+ males with 20 WT Benakeion females in biological triplicates following a standardised lab procedure (31).

### Molecular characterization of F1 intersex and male flies

gDNA from DsRed+/GFP+ F1 flies from Cas9-expressing females (DmPub.2,4; CcPub.1) crosses with gRNA-expressing males (GuideA.1-2; GuideB.1-2) was extracted using an adapted Chelex 100 Resin (Bio-Rad) protocol. In short, individual flies were incubated with resuspended Chelex 100 Resin (Bio-Rad) and Proteinase K (Thermo Fisher Scientific Inc.), followed by a precipitation with sodium acetate (3M) (Thermo Fisher Scientific Inc.) and ethanol absolute (VWR Life Science). gDNA was used as template for *ß-tub* homologue and tra target PCRs set up with Phusion High-Fidelity PCR Master Mix with HF Buffer (New England Biolabs® Inc.) using tra_F/tra_R and btub_F/btub_R primer pairs (Supplementary Table 6). PCR products were subsequently purified, and Sanger sequenced (Genewiz Inc.). The Sanger sequences were analysed using the DECODR v3.0 software (65) to estimate cleavage rates and predict mutation types. To distinguish XX and XY individuals, a karyotyping PCR was performed on gDNA with DreamTaq PCR Master Mix (2X) (Thermo Fisher Scientific Inc.) and CcYF/CcYR primers (66) (Supplementary Table 6).

### Modelling of Ceratitis capitata SIT

Individual-based simulations were generated with the forward genetic simulation software SLiM (version 4.2) (67). In this model, we incorporated medfly lifespan and demographic factors, which progress in weekly time steps, allowing for overlapping generations. Transgenic males are released in varying numbers each week after first allowing the simulation to equilibrate for 50 weeks. We assumed that the released transgenic males are sterile, causing females that mate with them to produce no offspring.

Typically, medflies have a 20% to 50% likelihood of mating more than once in their lifetime, with a preference for the first male they select (68). In our model based on adult mortality, we incorporated a 1/3 weekly remating probability for females. Additionally, sperm from the first mated male is preferred over sperm from subsequent matings (69). We assumed conservatively that if a female mates with a SIT male initially, a subsequent mating with a wild-type male allows for 50% fertility (50% chance to produce offspring if it would normally take place). Conversely, if the first mating is with a wild-type male, subsequent mating with a SIT male has no negative effect on fertility. At the beginning of each week, mating takes place between adults. Males are randomly selected during mating at a rate proportional to their fitness. pgSIT males are assumed to have no fitness costs, but radiation-based SIT males have a relative fitness of 0.83 based on mating ratios in an experimental study (58).

For mated females each week (except the first week as an adult) (70), offspring production takes place. The number of offspring is generated using a Poisson distribution with a mean of 9.37, based on the number of eggs laid per week by medflies (71). The survival rate of medfly larvae is strongly influenced by clutch density due to resource competition (72). Consequently, population density directly impacts larval survival. The density-based survival rate is based on a Beverton-Holt model and is calculated using the formula:*survival rate* = 0.3344/9.373 ∗ *β*/(*β* − 1 ∗ *f* + 1), where *β* represents the low-density growth rate (10 as default), and *f* denotes the competition factor, defined as the ratio between the actual number of larvae and the expected number of larvae. The constant is set to produce a stable population size at carrying capacity in the absence of SIT males. After one week, the juveniles enter the pupal stage for two weeks, after which they emerge as adults that do not require additional resources. Therefore, no density-dependent mortality occurs in the model during these stages, but baseline survival takes mortality of these stages into account.

Medfly males exhibit a longer lifespan compared to females. In our model, females do not survive beyond their thirteenth week, while males can live up to nineteen weeks. Both sexes experience a 100% survival rate during the first five weeks (juvenile stages, where all mortality is represented at the larval competition stage), after which age-based survival declines (producing a linear decline in number of surviving individuals), reaching 0% at week 19 for males and week 13 for females, based on a study examining lifespan across several strains (71). We set the age-based survival rate for each individual to be [1,1,1,1,1,13/14,12/13,11/ 12,10/11,9/10,8/9,7/8,6/7,5/6,4/5,3/4,2/3,1/2,0]for males and [1,1,1,1,1,7/8,6/7,5/ 6,4/5,3/4,2/3,1/2,0] for females. In reality, adult mortality can result from various other factors such as predation, but the level of this had not been evaluated by field studies. To account for this, we introduced a general mortality rate parameter that applies to adults of all ages, with a default value set at 0.9. The was set to produce a significant source of mortality, but age-based mortality remains important as well.

Sterile males were released at varying ratios, referring to release effort per week as a fraction of the number of wild-type males when the population is at carrying capacity. It directly represents the number of males for classic SIT, but the release for pgSIT males can be different based on relative juvenile mortality (which can reduce the actual release number) and fraction of fertile XX males (which can increase the actual release number). The simulation begins with an initial population of 50,000 adult females and is concluded 100 weeks after the SIT release, unless the population is eliminated earlier. All SLiM scripts and raw data can be accessed at https://github.com/jchamper/Medfly-SIT. The simulations were conducted using the High-Performance Computing Platform at the Center for Life Sciences, Peking University.

### Figure construction and statistical analyses

RStudio and Python were used for all statistical analysis and statistical significance was determined at *p*-value of *p* < 0.05. Graphs were similarly constructed in RStudio/Python, fly diagrams were created using Inkscape 1.3.2 (73), whilst all other figures were assembled in Microsoft PowerPoint.

## Data availability

Source data contains the data used for figure generation. Plasmids are available at addgene.org (#226752- 226757). The herein generated *C. capitata* transgenic strains are available upon request to A.M.

## Author contributions

A.M, O.S.A and E.B conceived the project and directed the research. J.L and N.P.K designed constructs and J.L performed the cloning. A.M and S.D designed the experiments. A.M performed the germline transformation. S.D, J.M and K.P maintained the strains. S.D performed the medfly and molecular validation experiments with help from K.P. Y.L and J.C generated the model. A.M and S.D analysed the data. All authors contributed to writing and editing of the manuscript and approved the final article.

## Acknowledgements

This work was supported by cooperative agreements (23-8130-1007-IA and AP23PPQS&T00C108) between the United States Department of Agriculture (USDA)— Animal and Plant Health Inspection Service (APHIS)— Plant Protection and Quarantine (PPQ) and Imperial College London and University of California—San Diego, respectively. Mention of trade names or commercial products in this publication is solely for the purpose of providing specific information and does not imply recommendation or endorsement by the U.S. Department of Agriculture, an equal opportunity employer. J.M. is supported by funding that was provided by the European Union’s Horizon Europe Research and Innovation Programme REACT to A.M. (Grant agreement 101059523).

## Supplementary Information legends

**Supplementary Figure 1.**
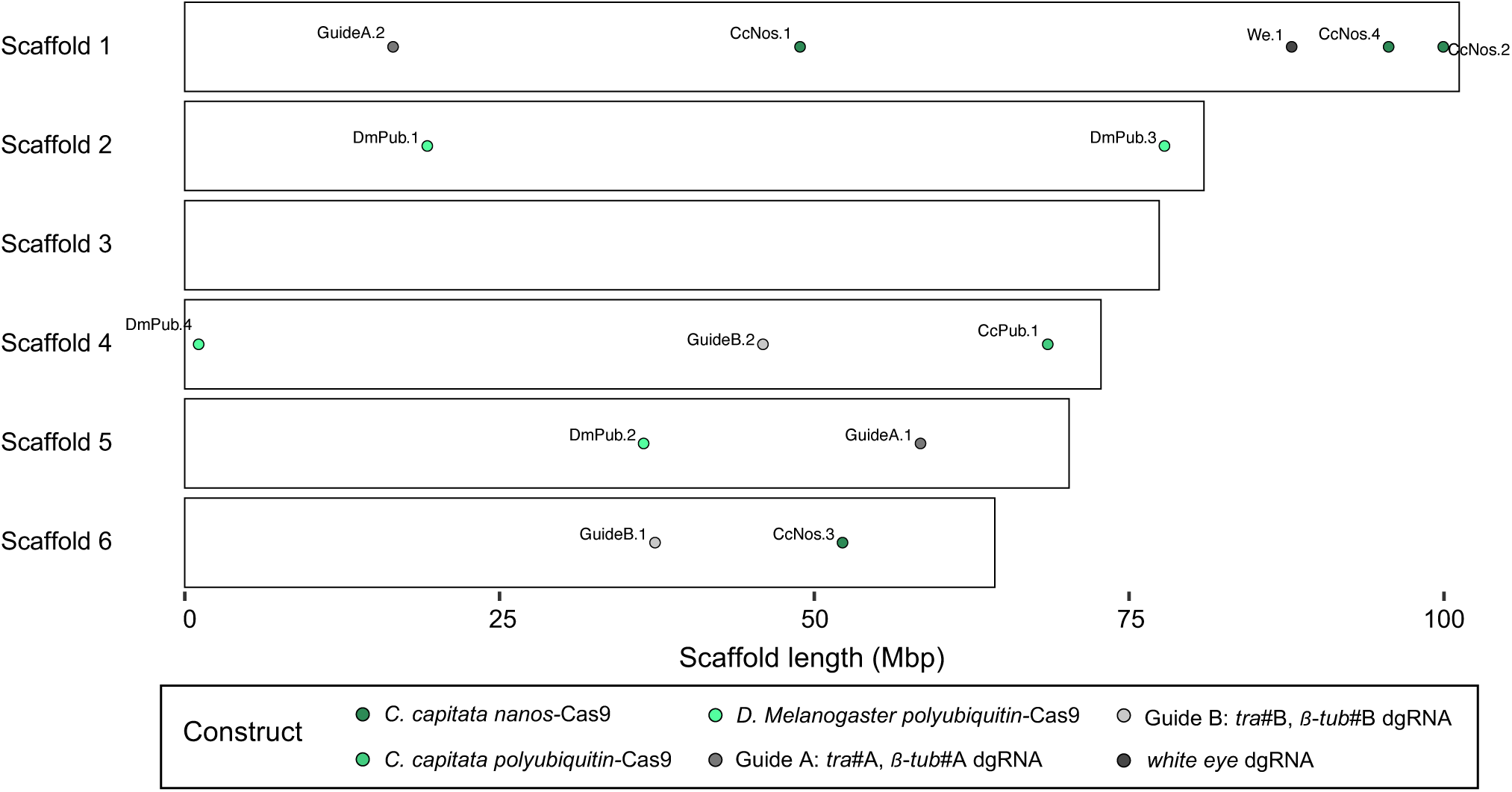
Integrations of *piggyBac* constructs. The map depicting the genomic integrations for the 3 Cas9 and 3 double-guide RNA (dgRNA) *piggyBac* constructs used in this study. As a result of germline transformation, 14 strains with unique integrations were established and characterised via inverse PCR using the GenBank GCA_905071925.1 *C. capitata* genome re-assembly. Constructed in RStudio.

**Supplementary Figure 2.**
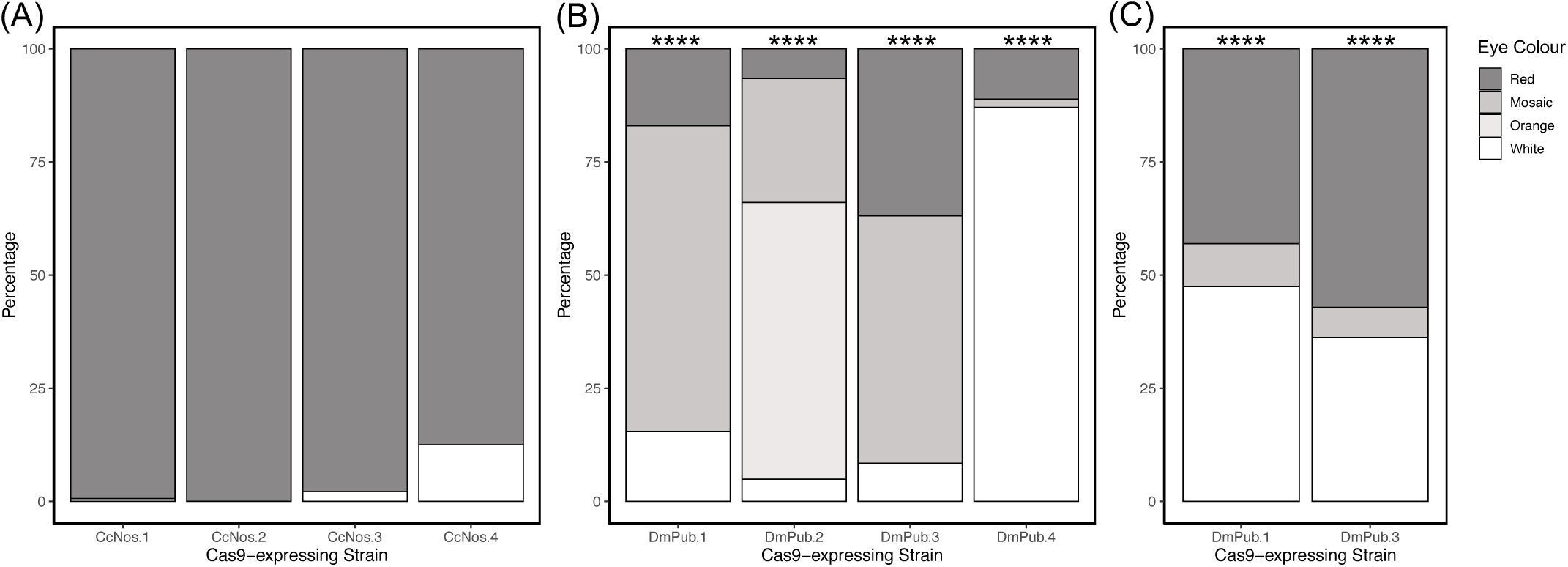
Optimisation of the CRISPR/Cas9 toolkit against the *white eye* gene. Stack graphs showing eye colour percentages of (A) *Ceratitis capitata nanos*-Cas9 F2, (B) *Drosophila melanogaster polyubiquitin*-Cas9 F1, and (C) *D. melanogaster polyubiquitin*-Cas9 F2 progeny. Individuals were exclusively screened for eye colour and scored as ‘red’, ‘mosaic’, ‘orange’ or ‘white’. (B) F1 population is the direct result of crosses between Cas9-harbouring females with dgRNA-harbouring (We.1) males. (A, C) F2 populations are a result of crosses between DsRed+/GFP+ red-eyed F1 females and males from the homozygous *white eye* mutant (-/-) strain. Significance levels for the chi-squared test are indicated as follows: *p* < 0.05 = *, *p* < 0.01 = **, *p* < 0.001 = *** and *p* < 0.0001 = ****. Constructed in RStudio.

**Supplementary Figure 3.**
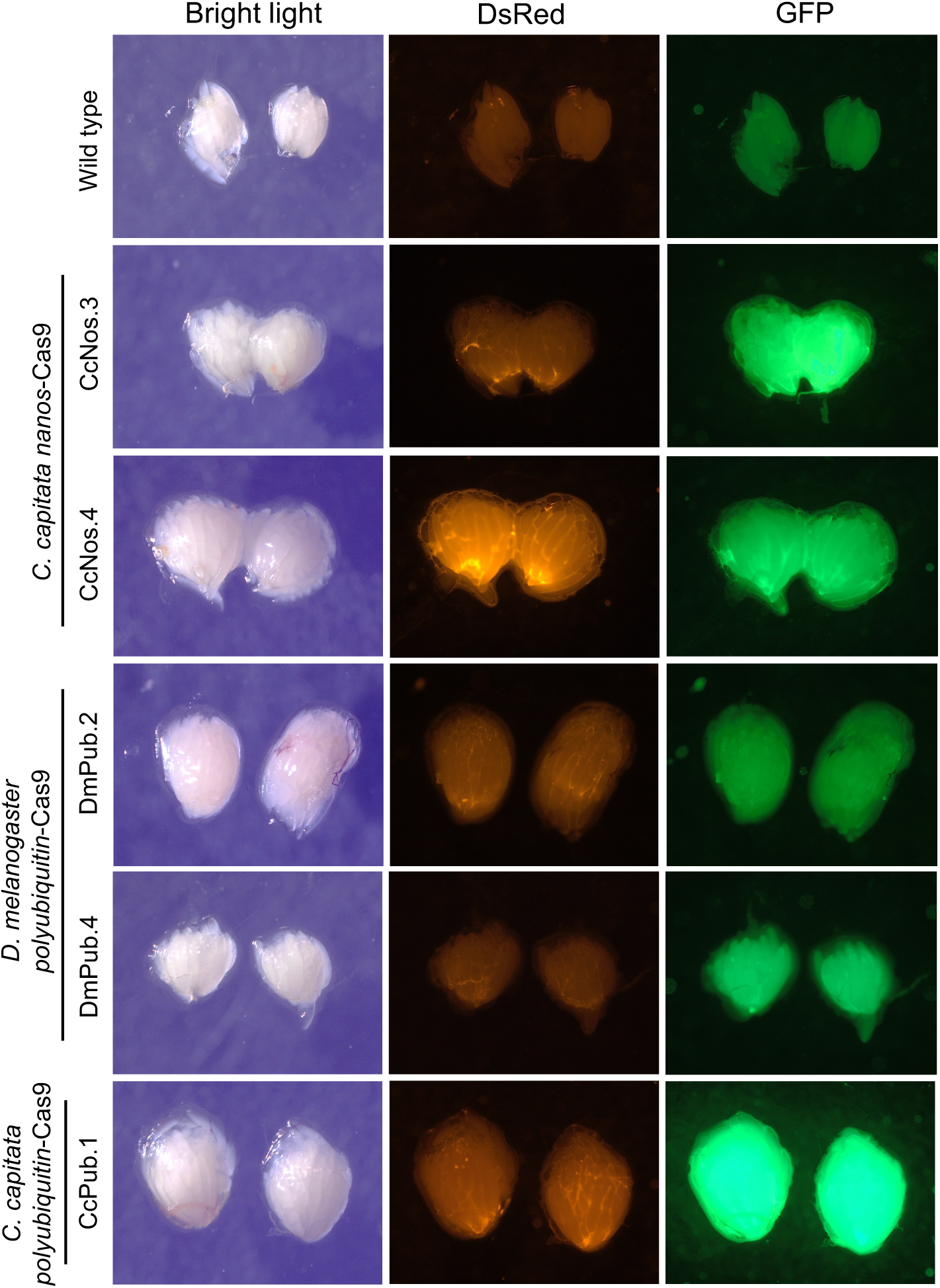
Fluorescence of Cas9-harbouring female ovaries. Representative ovaries, dissected from sexually mature females, were imaged under red fluorescence protein (RFP) and green fluorescence protein (GFP) filters, in addition to bright light with standardised settings. All Cas9 cassettes possess a constitutively expressed DsRed transformation marker and are equipped with GFP, which is linked to the upstream Cas9 via a T2A peptide. Dissections were performed on homozygous females from CcNos.3, CcNos.4, DmPub.2 and CcPub.1 strains, whilst for DmPub.4 heterozygous females were dissected instead due to a lack of homozygous individuals in the strain.

**Supplementary Figure 4.**
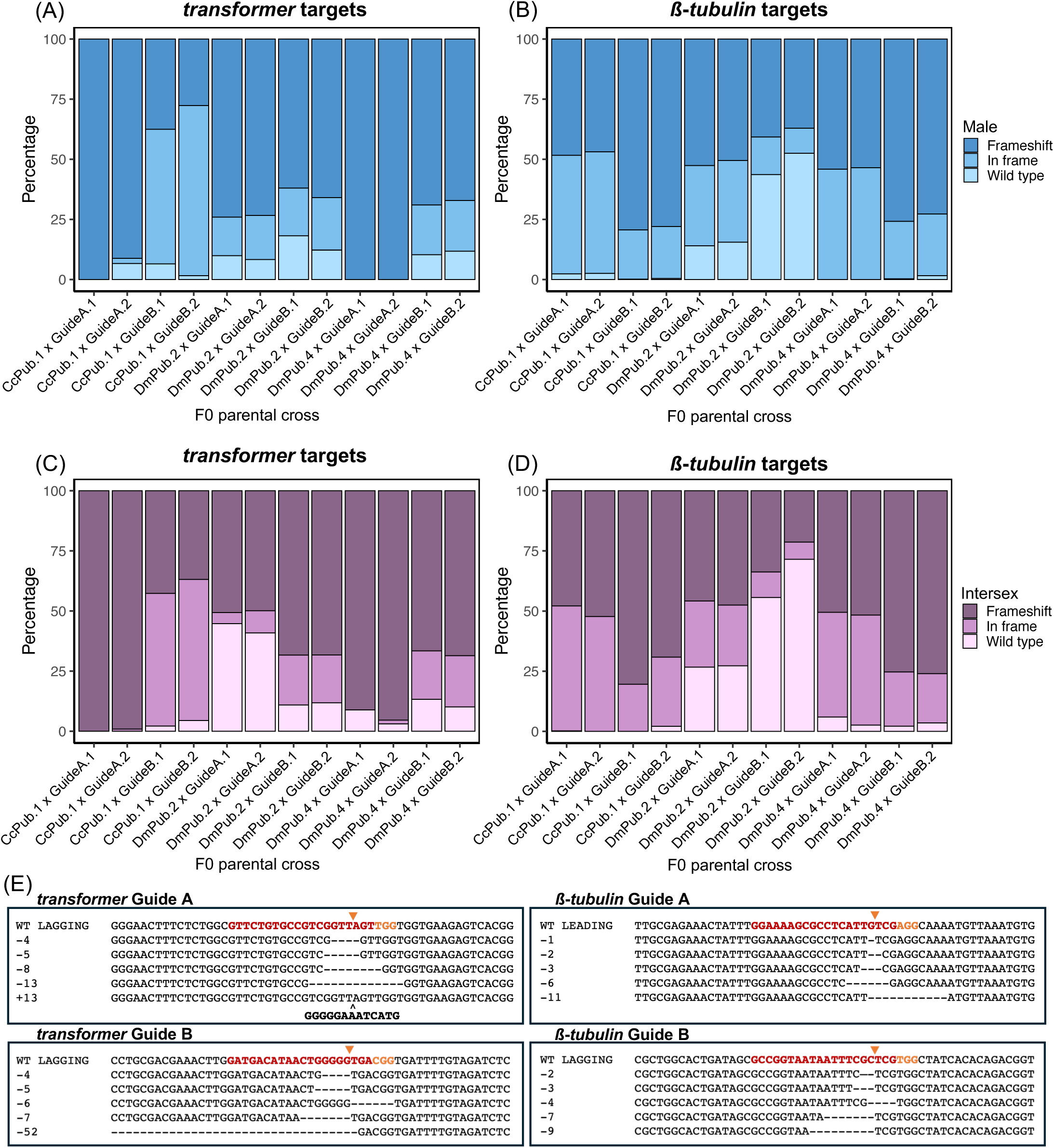
Characterisation of gRNA efficiency in F1 trans-heterozygotes. (A-D) Stack graphs showing mean predicted sequence identities for (A, C) *transformer* and (B, D) *ß-tubulin* gRNA targets in DsRed+/GFP+ F1 (A, B) males and (C, D) intersexes. (A-D) The averages were obtained from 3-11 sequenced males or intersexes per F0 cross. DECODR v3.0 (65) was used for sequence deconvolution whereby individual sequence identities were broken down into wild-type, in-frame indel or frameshift by percentage. (E) Example indels identified through sequence deconvolution for the 4 used gRNAs, whereby sequences are shown with 5’-3’ gRNA direction.

**Supplementary Figure 5.**
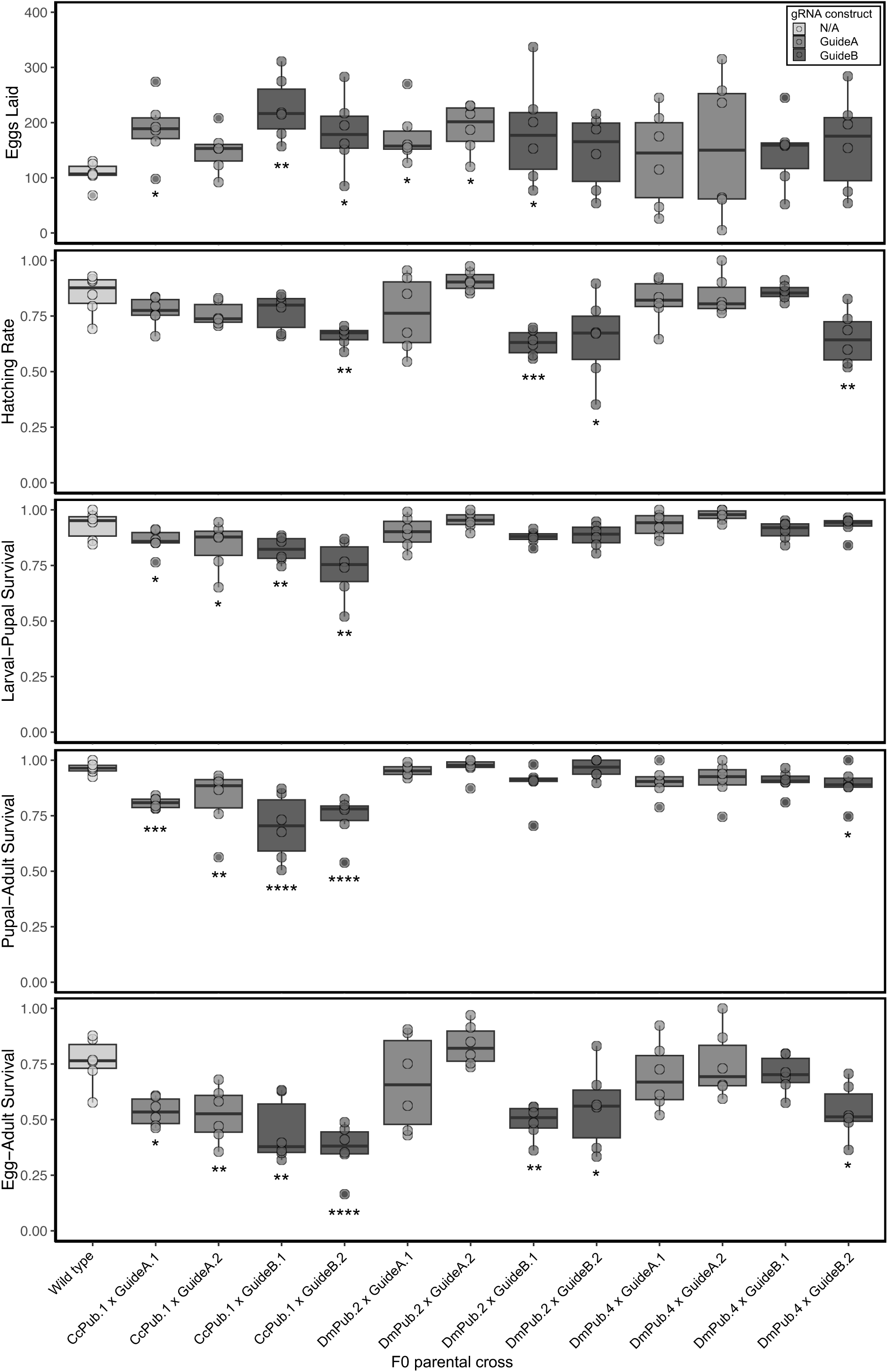
Fitness of F1 trans-heterozygotes. Boxplots showing egg-adult survival of F1 trans-heterozygotes compared to wild-type controls measured via eggs laid with corresponding hatching rates, hatched larval-pupal, pupal-adult and total egg-adult recovery rates. Total eggs assessed were collected within a 5-hour period from 6 replicate crosses between 5 dgRNA-harbouring males and 15 Cas9-harbouring females. The boxes represent the interquartile ranges, the lines represent the mean values, the whiskers represent minima and maxima, and the dots represent raw replicate values with outliers of 1.5 times interquartile range found outside the whiskers. The statistically significant wild-type – transgenic differences are displayed on the bar charts as follows: *p* < 0.05 = *, *p* < 0.01 = **, *p* < 0.001 = *** and *p* < 0.0001 = **** (Dunn’s test). Constructed in RStudio.

**Supplementary Figure 6.**
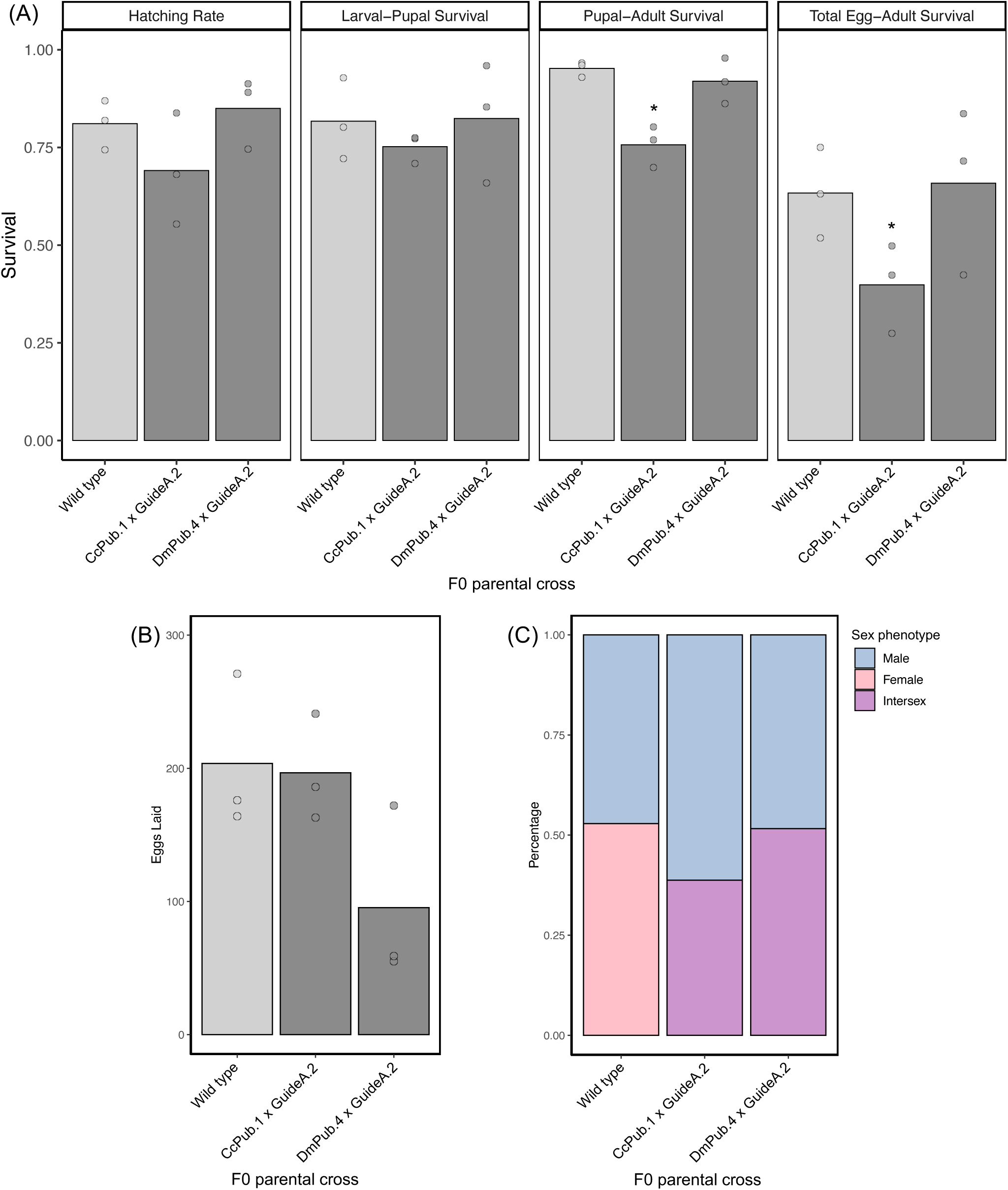
Fitness of F1 trans-heterozygotes with varied sex-conversion rates. (A) Bar charts showing egg-adult survival of F1 trans-heterozygotes compared to wild-type controls showing hatching rates, hatched larval-pupal, pupal-adult and total egg-adult recovery rates. (B) A bar chart showing corresponding eggs laid, collected within a 5-hour period from triplicates crosses between 10 dgRNA-harbouring males and 20 Cas9-harbouring females. (C) A stack graph showing the corresponding mean adult phenotypes which were scored as ‘male’, ‘intersex’ or ‘female’. Flies were screened for DsRed and GFP and all non-wild-type F1 progeny was DsRed+/GFP+. (A-B) The bar levels represent the mean values, and the dots represent raw replicate values. The statistically significant wild-type – transgenic differences are displayed on the bar charts as follows: *p* < 0.05 = * (Dunn’s test). Constructed in RStudio.

**Supplementary Figure 7.**
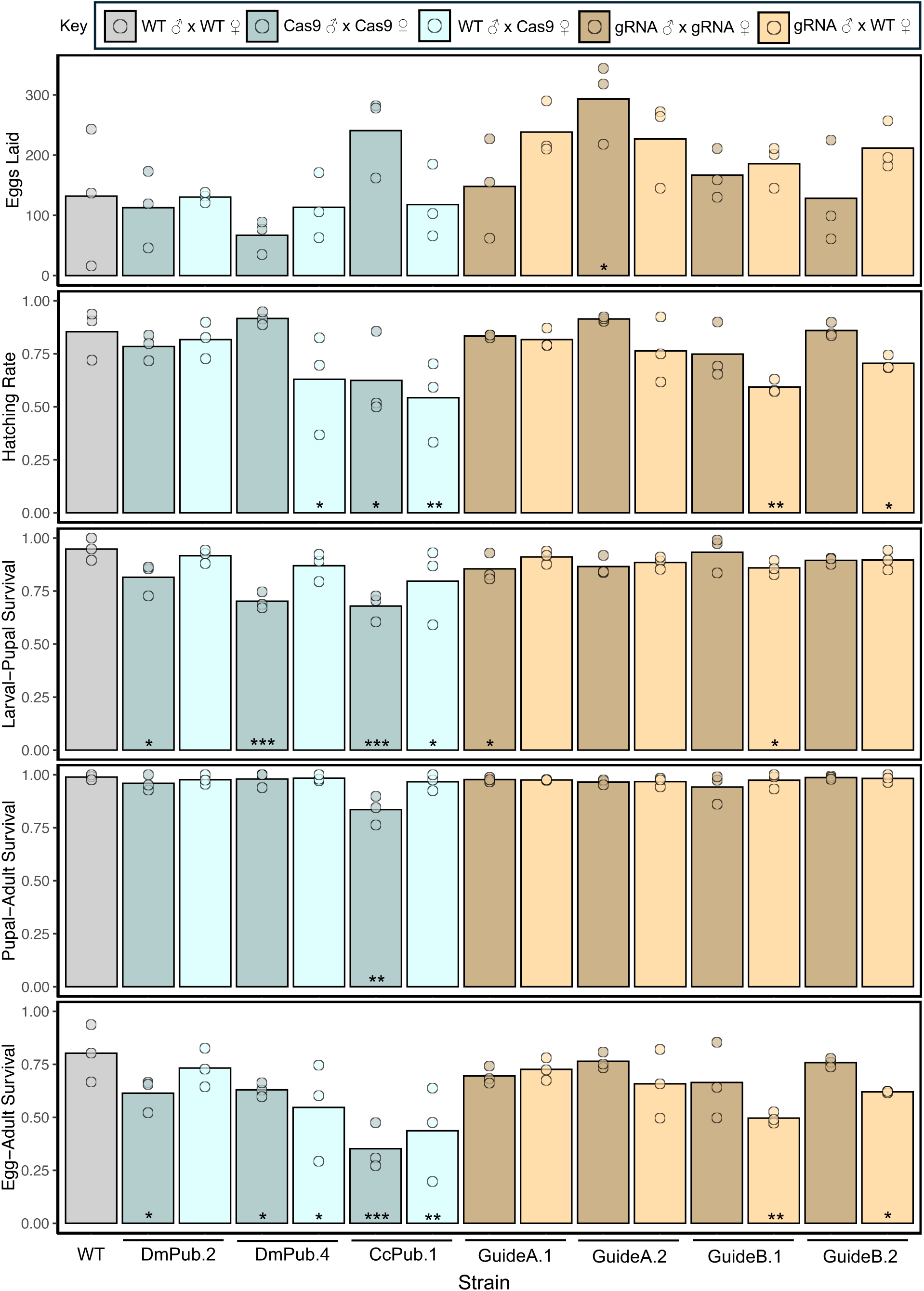
Fitness of *piggyBac* Cas9 and dgRNA-harbouring strains. Bar charts showing transgenic strain egg-adult survival compared to wild-type controls measured via eggs laid with corresponding hatching rates, hatched larval-pupal, pupal-adult and total egg-adult recovery rates. Total eggs assessed were collected within a 5-hour period from triplicate crosses between 10 males and 20 females. Each Cas9-harbouring strain (DmPub.2, DmPub.4, CcPub.1), depicted in blue, was subjected to sibling crosses and crosses of transgenic females with wild-type males. Each dgRNA-harbouring strain (GuideA.1-2; GuideB.1-2), depicted in beige, was subjected to sibling crosses and crosses of transgenic males with wild-type females. The bar levels represent the mean values, and the dots represent raw replicate values. The crosses performed are additionally summarised in Supplementary Table 5. The statistically significant wild-type – transgenic differences are displayed on the bar charts as follows: *p* < 0.05 = *, *p* < 0.01 = ** and *p* < 0.001 = *** (Dunn’s test). Constructed in RStudio.

**Supplementary Figure 8.**
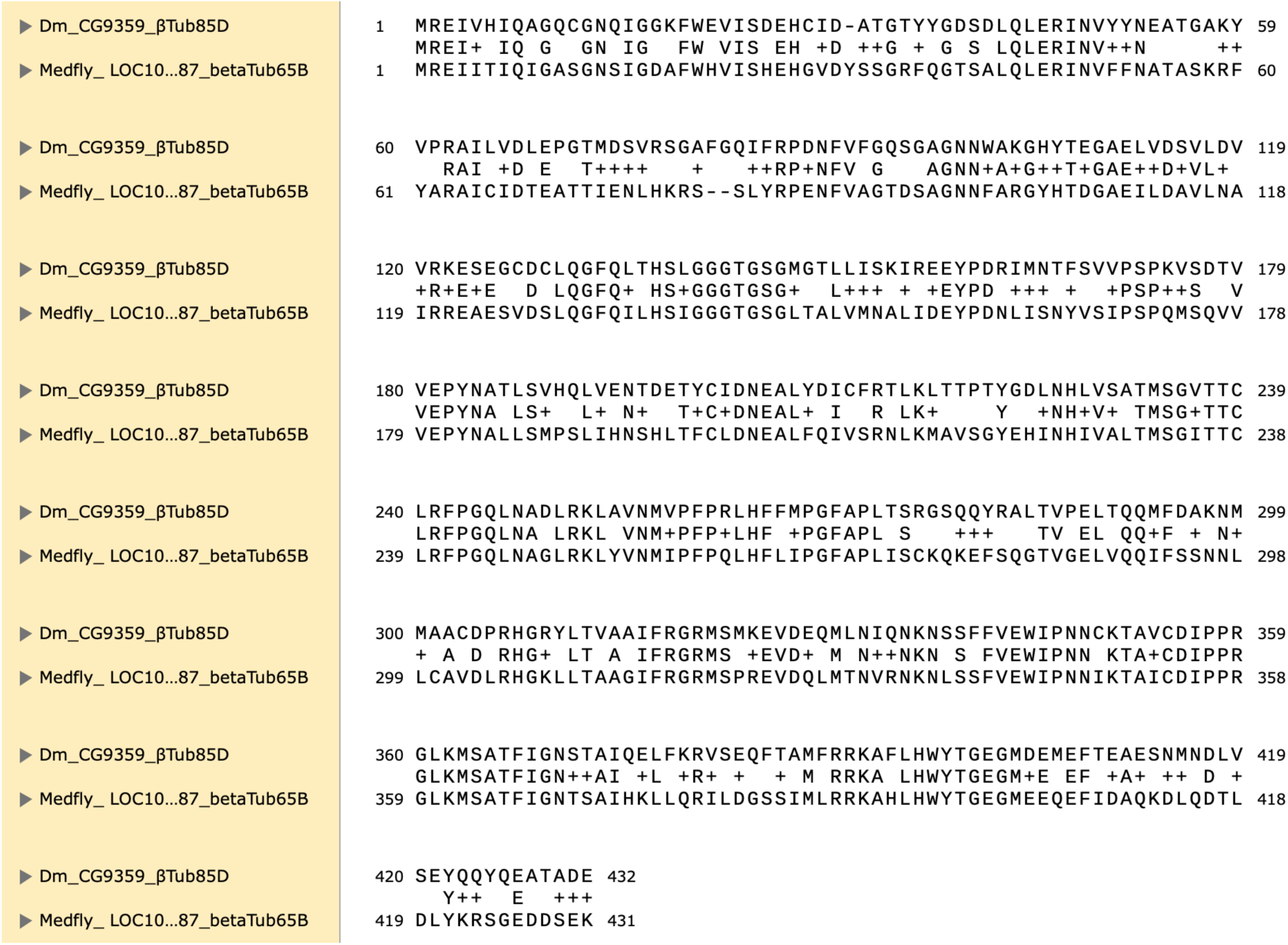
**Protein alignment of *ß*-*tubulin in Drosophila melanogaster* and *Ceratitis capitata* showed a 73.90% similarity**

**Supplementary Figure 9.**
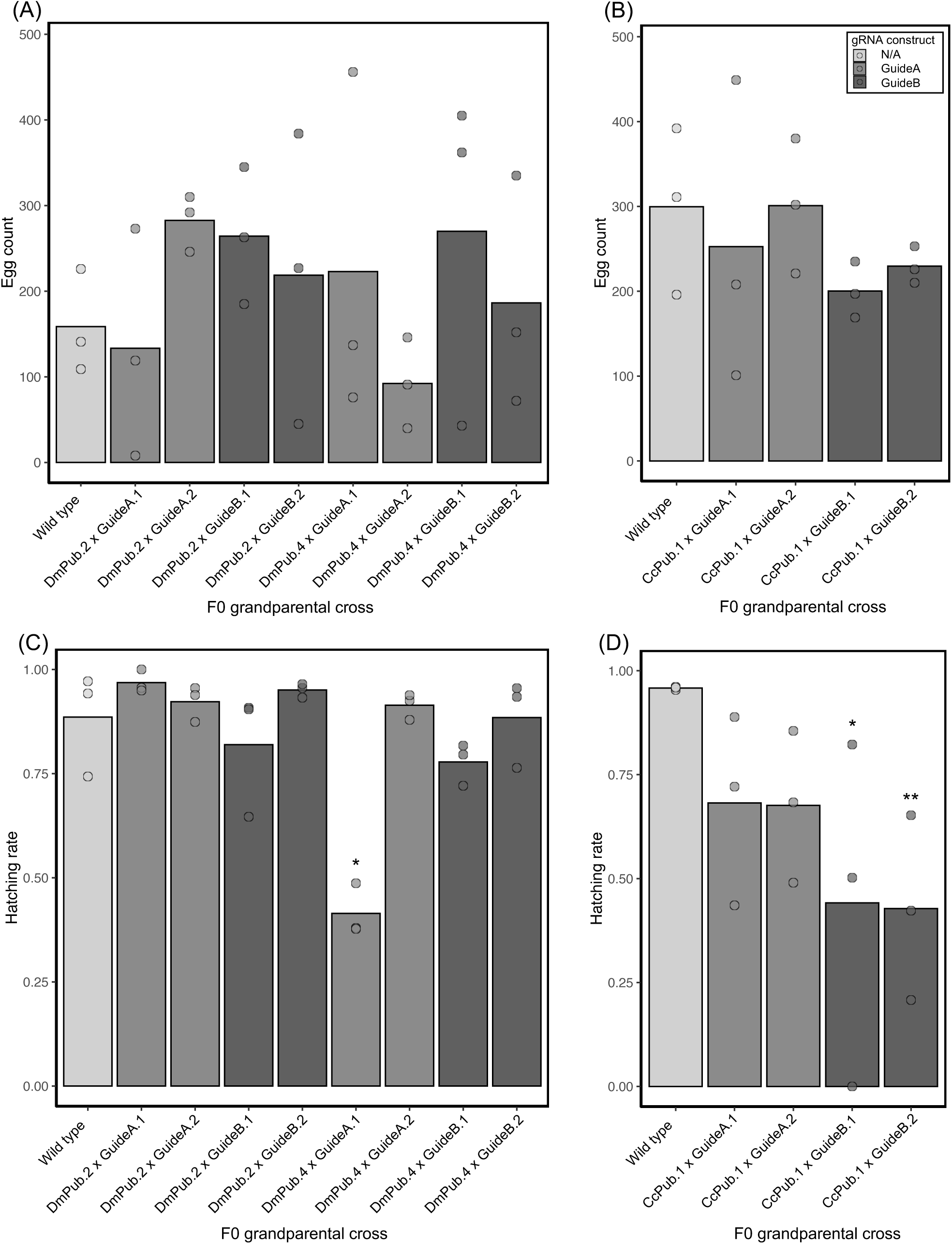
Characterisation of trans-heterozygous F1 males. Bar charts showing egg laying (A, B) and corresponding egg hatching (C, D) rate comparisons between wild-type sibling crosses with DsRed+/GFP+ F1 males crossed with wild-type females. Egg laying and egg hatching were measured separately for the wild-type-crossed F1 males from (A, C) F0 *Drosophila melanogaster polyubiquitin*-Cas9 (DmPub.1-4)-harbouring females with the dgRNA (GuideA.1-2; GuideB.1-2)-harbouring males; and (B, D) F0 *Ceratitis capitata polyubiquitin*-Cas9 (CcPub.1)-harbouring females with the dgRNA (GuideA.1-2; GuideB.1-2)-harbouring males. (A-D) Total eggs assessed were collected within a 5-hour period from triplicate crosses between 10 males and 20 females. The bar levels represent the mean values, and the dots represent raw replicate values. The statistically significant wild-type – transgenic differences are displayed on the bar charts as follows: *p* < 0.05 = * and *p* < 0.01 = ** (Dunn’s test). Constructed in RStudio.

**Supplementary Table 1.**
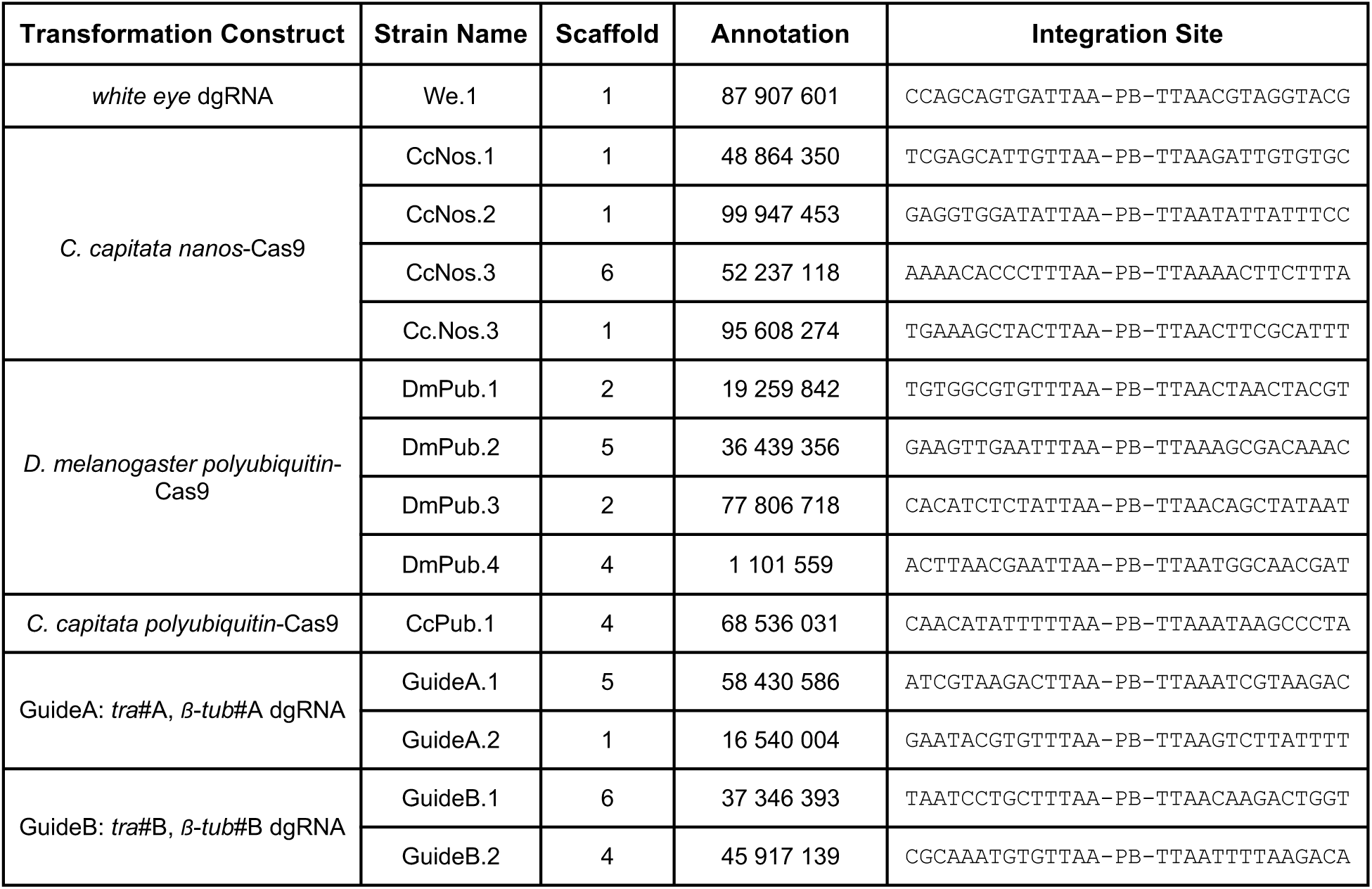
Genomic integration annotations of *piggyBac* constructs. The genomic sequences flanking the *piggyBac* constructs as determined through inverse PCR and visually represented in Supplementary Figure 1. Altogether 14 strains with unique integrations were established in this study.

**Supplementary Table 2.** Eye colour phenotypes of DsRed-/GFP+ F1 progeny. The Cas9-harbouring females were crossed with males from the *white eye*-dgRNA-harbouring We.1 strain. Their progeny was screened for fluorescence and classified by eye colour thereafter.

|  | Eye colour |  |  |  |
| --- | --- | --- | --- | --- |
| Cross | Red | Mosaic | Orange | White |
| DmPub.2 x We.1 | 7 |  |  |  |
| DmPub.4 x We.1 |  | 4 |  | 147 |
| CcPub.1 x We.1 |  |  |  | 106 |

**Supplementary Table 3.** Sex phenotypes of DsRed-/GFP+ F1 progeny. The Cas9-harbouring and dgRNA-harbouring strains were reciprocally crossed together, and their progeny was scored as ‘Male’, ‘Intersex’ or ‘Female’ post fluorescence screening.

|  |  | Sex phenotype |  |  |
| --- | --- | --- | --- | --- |
| Source of Cas9 | Cross | Female | Intersex | Male |
| Maternal | DmPub.2 x GuideA.1 | 1 |  | 1 |
|  | DmPub.2 x GuideA.2 |  |  | 1 |
|  | DmPub.2 x GuideB.1 |  |  |  |
|  | DmPub.2 x GuideB.2 |  |  |  |
|  | DmPub.4 x GuideA.1 |  | 47 | 159 |
|  | DmPub.4 x GuideA.2 |  | 76 | 285 |
|  | DmPub.4 x GuideB.1 |  | 63 | 96 |
|  | DmPub.4 x GuideB.2 |  | 135 | 144 |
|  | CcPub.1 x GuideA.1 |  | 1 | 7 |
|  | CcPub.1 x GuideA.2 |  | 5 | 28 |
|  | CcPub.1 x GuideB.1 |  | 51 | 60 |
|  | CcPub.1 x GuideB.2 |  | 73 | 82 |
| Paternal | DmPub.2 x GuideA.1 |  |  |  |
|  | DmPub.2 x GuideA.2 |  |  |  |
|  | DmPub.2 x GuideB.1 |  |  |  |
|  | DmPub.2 x GuideB.2 |  |  |  |
|  | DmPub.4 x GuideA.1 | 58 |  | 51 |
|  | DmPub.4 x GuideA.2 | 80 |  | 92 |
|  | DmPub.4 x GuideB.1 | 45 |  | 43 |
|  | DmPub.4 x GuideB.2 | 97 |  | 112 |
|  | CcPub.1 x GuideA.1 | 24 |  | 24 |
|  | CcPub.1 x GuideA.2 | 20 |  | 17 |
|  | CcPub.1 x GuideB.1 | 22 |  | 28 |
|  | CcPub.1 x GuideB.2 | 9 |  | 12 |

**Supplementary Table 4.** Morphology of F1 intersex internal genitalia. Trans-heterozygous intersexes were abdominally dissected upon sexual maturation.

| Parental crosses | 2 full ovaries | 2 malformed ovaries | 1 full ovary | 1 malformed ovary | Nothing | 1 malformed testis | 1 full testis | 2 malformed testes | 2 full testes | Total |
| --- | --- | --- | --- | --- | --- | --- | --- | --- | --- | --- |
| GuideA x DmPub.2 | 2 | 2 |  |  |  |  |  |  | 1 | 5 |
| GuideA x DmPub.4 |  |  |  | 1 | 2 | 1 |  | 1 |  | 5 |
| GuideA x CcPub.1 |  |  |  |  | 4 | 1 |  |  |  | 5 |
| GuideB x DmPub.2 | 5 |  |  |  |  |  |  |  |  | 5 |
| GuideB x DmPub.4 |  | 1 |  |  | 4 |  |  |  |  | 5 |
| GuideB x CcPub.1 |  |  | 3 |  | 2 |  |  |  |  | 5 |
| Total | 7 | 3 | 3 | 1 | 12 | 2 | 0 | 1 | 1 | 30 |

**Supplementary Table 5.** Crosses performed in line egg-adult assay. The parental crosses for line egg-adult assay were performed in triplicates for 3 Cas9-harbouring and 4 dgRNA-harbouring strains alongside wild type controls. Both sibling and transgenic-wild type crosses were performed to mimic 2 and 1 transgene copy inheritance, respectively.

| Construct type | Females | Males |
| --- | --- | --- |
| Cas9 | DmPub.2 homozygotes | DmPub.2 homozygotes |
|  | DmPub.2 homozygotes | Wild type |
|  | DmPub.4 heterozygotes | DmPub.4 heterozygotes |
|  | DmPub.4 heterozygotes | Wild type |
|  | CcPub.1 homozygotes | CcPub.1 homozygotes |
|  | CcPub.1 homozygotes | Wild type |
| dgRNA | GuideA.1 homozygotes | GuideA.1 homozygotes |
|  | Wild type | GuideA.1 homozygotes |
|  | GuideA.2 homozygotes | GuideA.2 homozygotes |
|  | Wild type | GuideA.2 homozygotes |
|  | GuideB.1 homozygotes | GuideB.1 homozygotes |
|  | Wild type | GuideB.1 homozygotes |
|  | GuideB.2 homozygotes | GuideB.2 homozygotes |
|  | Wild type | GuideB.2 homozygotes |
| N/A | Wild type | Wild type |

**Supplementary Table 6.**
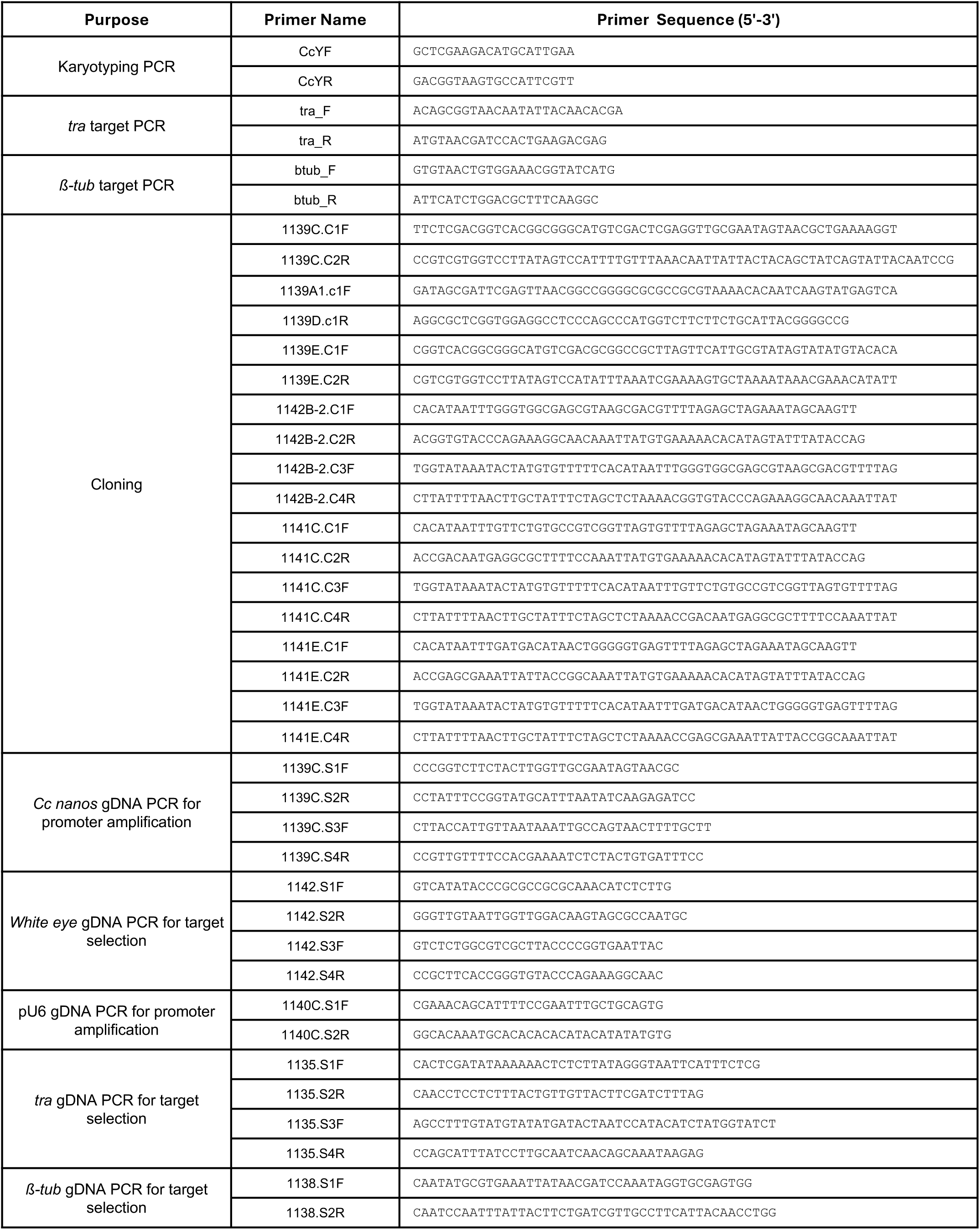
Primer summary. The primers used for F1 trans-heterozygote characterisation and plasmid cloning. The karyotyping PCR was performed with previously described primers (66).

